# IST1 regulates select recycling pathways

**DOI:** 10.1101/2023.07.31.551359

**Authors:** Amy K. Clippinger, Teresa V. Naismith, Wonjin Yoo, Silvia Jansen, David J. Kast, Phyllis I. Hanson

## Abstract

ESCRTs (Endosomal Sorting Complex Required for Transport) are a modular set of protein complexes with membrane remodeling activities that include the formation and release of intralumenal vesicles (ILVs) to generate multivesicular endosomes. While most of the 12 ESCRT-III proteins are known to play roles in ILV formation, IST1 has been associated with a wider range of endosomal remodeling events. Here, we extend previous studies of IST1 function in endosomal trafficking and confirm that IST1, along with its binding partner CHMP1B, contributes to scission of early endosomal carriers. Depleting IST1 impaired delivery of transferrin receptor from early/sorting endosomes to the endocytic recycling compartment and instead increased its rapid recycling to the plasma membrane via peripheral endosomes enriched in the clathrin adaptor AP-1. IST1 is also important for export of mannose 6-phosphate receptor from early/sorting endosomes. Examination of IST1 binding partners on endosomes revealed that IST1 interacts with the MIT domain-containing sorting nexin SNX15, a protein previously reported to regulate endosomal recycling. Our kinetic and spatial analyses establish that SNX15 and IST1 occupy a clathrin-containing subdomain on the endosomal perimeter distinct from those previously implicated in cargo retrieval or degradation. Using live-cell microscopy we see that SNX15 and CHMP1B alternately recruit IST1 to this subdomain or the base of endosomal tubules. These findings indicate that IST1 contributes to a subset of recycling pathways from the early/sorting endosome.

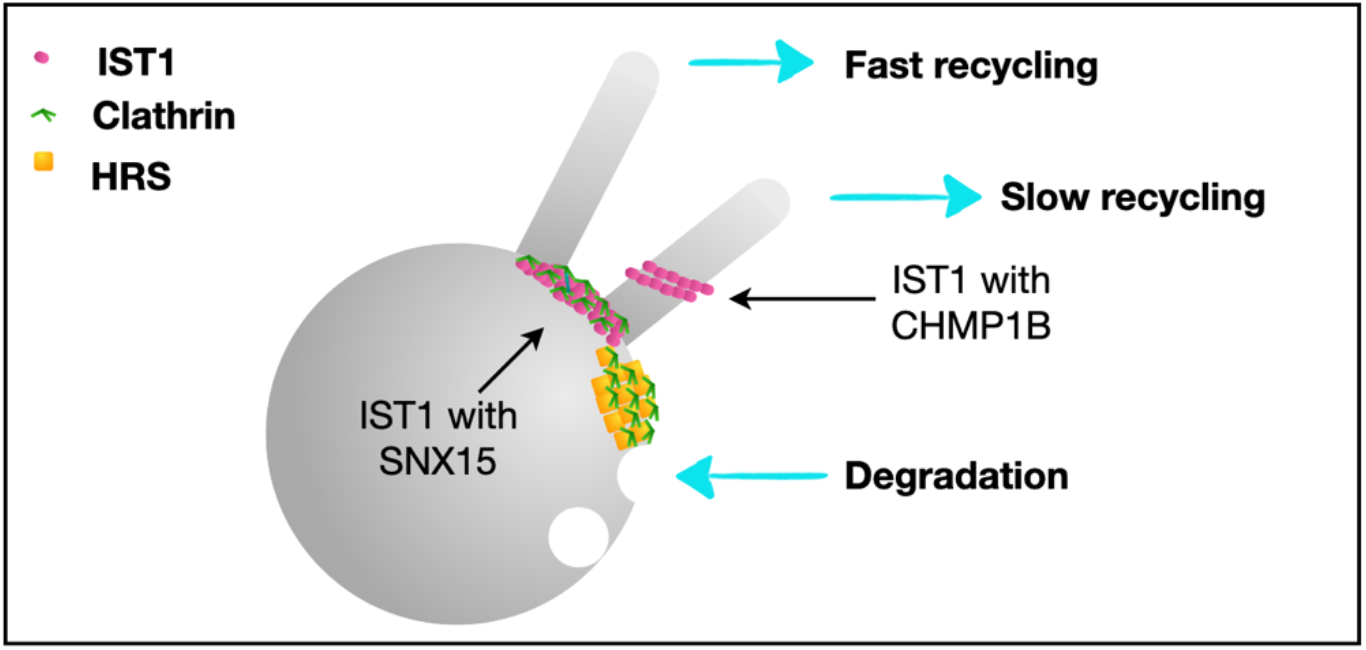

## Introduction

ESCRT (Endosomal Sorting Complex Required for Transport)-III proteins are primarily cytosolic monomers that assemble to form membrane-associated filamentous polymers which mediate membrane remodeling events including intraluminal vesicle (ILV) formation, virus budding, nuclear envelope closure, and cytokinetic abscission (Vietri *et al*., 2020). ESCRT-III proteins contain a “core” ESCRT-III domain that enables homo-and heteropolymer assembly followed by C terminal MIM motifs that interact with MIT domain containing proteins. IST1 is structurally and functionally distinct from other ESCRT-III proteins. Its “core” ESCRT-III domain only co-assembles with CHMP1A/1B (Bajorek *et al*., 2009a). IST1 also has not one, but two distinct MIM domains, as well as an extended unstructured and proline-rich linker region between its “core” and MIM domains. IST1 interacts via these domains with general ESCRT-III as well as numerous IST1-specific partners (Yang *et al*., 2008; Renvoisé *et al*., 2010; Allison *et al*., 2013, 2017; Maemoto *et al*., 2014; Chang *et al*., 2019). Among its best-studied interactors is Vps4 (VPS4A/4B in mammalian cells), the AAA ATPase responsible for disassembling ESCRT-III polymers (Saksena *et al*., 2009; Adell *et al*., 2014; Mierzwa *et al*., 2017). Interestingly, VPS4A mutations that specifically impede release of IST1, but not another ESCRT-III protein CHMP2B, from membranes result in abnormal neurodevelopment (Rodger *et al*., 2020). Mutations in IST1 specific interactors spastin (SPAST) and spartin (SPART) have also been directly linked to hereditary spastic paraplegia (HSP) (Hazan *et al*., 1999; Patel *et al*., 2002; Blackstone *et al*., 2011).

As might be expected based on its unique structure compared to other ESCRT-III proteins, IST1 has been found to have diverse functional roles in endosomal trafficking across a variety of systems. ESCRT-IIIs, similar to the rest of the ESCRT machinery, are responsible for generating intraluminal vesicles (ILVs) within multivesicular bodies (MVBs) to enable lysosomal degradation and were discovered in a now classic screen for vacuolar protein sorting (vps) mutants (Odorizzi *et al*., 1998; Babst *et al*., 2002; Teis *et al*., 2008). Notably, IST1 was not part of the original set of ESCRTs and was instead later identified on the basis of genetic interactions with and structural homology to other ESCRT-III proteins (Rue *et al*., 2008; Bajorek *et al*., 2009a, 2009b). IST1 is not required for ILV formation in *Saccharomyces cerevisiae* (Dimaano *et al*., 2008; Rue *et al*., 2008; Xiao *et al*., 2009; Nickerson *et al*., 2010), plants (Buono *et al*., 2016), or HeLa cells (Allison *et al*., 2013). Instead, it has been found to function in endosomal recycling and retrograde trafficking pathways (Allison *et al*., 2013, 2017, 2019; Laidlaw *et al*., 2022a). Indeed, its unique assembly with CHMP1B creates constricting filaments around (instead of within) membrane tubules (McCullough *et al*., 2015) with proposed roles in tubule scission (Nguyen *et al*., 2020; Cada *et al*., 2022). However, IST1 is also both a negative and positive regulator of Vps4 activity (Tan *et al*., 2015) and is a positive modulator of MVB biogenesis in yeast (Dimaano *et al*., 2008; Rue *et al*., 2008; Xiao *et al*., 2009; Nickerson *et al*., 2010). It also controls overall ESCRT recruitment and as a consequence ILV formation during early oocyte maturation in *C. elegans* (Frankel *et al*., 2017). Altogether, IST1 appears to be a multifunctional ESCRT-III protein.

To gain further insight into how IST1 affects endosomal dynamics, we set out to learn more about where and how it functions in endosomal trafficking in cultured human cells. We find that IST1 is enriched on early/sorting endosomes where it localizes with its binding partners SNX15 or CHMP1B to endosomal subdomains that are distinct from those previously known to direct cargo for retrieval/recycling or degradation. Functionally, we find that IST1 is important for slow but not fast transferrin receptor recycling with a key role in delivering cargo from early/sorting to recycling endosomes. Our results provide insight into the diversity of machinery that controls exit from the early/sorting endosome and thereby cellular membrane organization.

## Results

### IST1 contributes to scission of tubules on early/sorting endosomes

To assess IST1’s contributions to endosomal homeostasis in mammalian cells, we asked where and how it functions in two human cell lines, T24 bladder carcinoma and U2OS osteosarcoma cells. An antibody recognizing endogenous IST1 confirmed its localization throughout the cytoplasm, at cell bridges and mid-bodies and on bright punctate structures that based on previous studies are likely to be early/sorting endosomes (Figure 1A) (Agromayor *et al*., 2009; Allison *et al*., 2017). Indeed, many of the IST1 marked structures were also immunoreactive for the early endosomal protein EEA1 (Figure 1B, Supplemental Figure S1).

**Figure 1.**
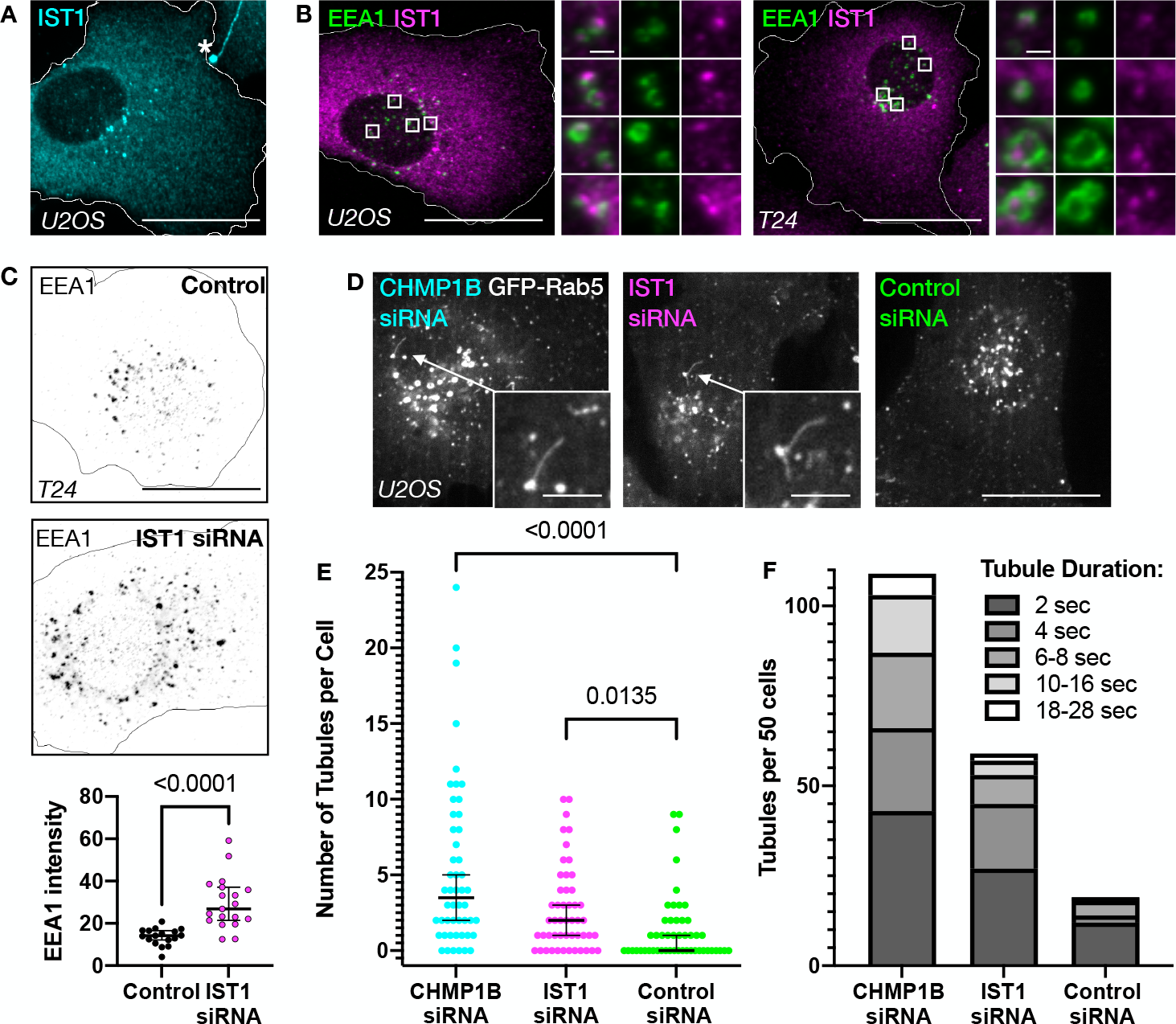
IST1 assembled with SNX15 but not CHMP1B strongly associates with vacuolar clathrin. **(A)** U2OS cells expressing SNX15GFP and mRFP CLCa. Inset panels show separate and merged channels of ROI.The relationship of SNX15 and clathrin on the vacuolar domain is denoted by arrow and on the tubular domain by arrow heads. Images are single slices. **(B)** U2OS cell expressing SNX15gfp and stained for AP-1, image is max intensity projection. The inset demonstrates collections of AP-1 puncta originating from SNX15gfp positive endosomes (empty arrow heads). **(C-E)** U2OS cells expressing IST1GFP/SNX15myc or IST1GFP/CHMP1B with mRFP CLCa. Images are single slices. The extent of colocalization between GFP tagged IST1 constructs and mRFP CLCa when assembled with SNX15myc or CHMP1B is quantified in (C). n=10 for each group. **(D and E)** Image sequences taken from the ROIs in cells showing in (C). **(F)** Model of IST1 localization to endosomal microdomains when co-assembled with SNX15 or CHMP1B, and IST1 function in endosomal recycling. Scale bars in the whole cells and/or insets are as follows: (A-B) 25 μm and 1 μm, respectively; (C-E) 25 μm and 0.5 μm respectively.

To explore IST1’s function, we depleted it using a previously validated siRNA duplex (Allison *et al*., 2013, 2017) that reduced IST1 by >90% in both T24 and U2OS cells (Supplemental Figure S2). By immunostaining, we found that cells depleted of IST1 had increased levels of endosomal EEA1 (Figure 1C) consistent with earlier reports of IST1 playing a role in the evolution of early/sorting endosomes via export of SNX1-and SNX-4 marked tubules (Allison *et al*., 2013). We also confirmed that a second IST1 siRNA duplex similarly decreased IST1 while increasing endosomal EEA1 (Supplemental Figure S2&S3). Previous studies of IST1’s role in endosomal recycling have attributed this to the partnership that we and others described between IST1 and its membrane-proximal binding partner CHMP1B on endosomal tubules (McCullough *et al*., 2015; Connell *et al*., 2019; Nguyen *et al*., 2020b; Cada *et al*., 2022). To visualize the dynamics of tubulovesicular carriers emerging from early/sorting endosomes, we monitored GFP-Rab5b by live-cell spinning disc confocal microscopy (Zeigerer *et al*., 2012; Rowland *et al*., 2014). GFP-Rab5b organelles are typically punctate and tubules that occasionally emerge are short-lived (Supplemental Figure S4). After depleting IST1 or CHMP1B, we measured tubule length and persistence and found that loss of either protein increased both tubule frequency and lifetime (Figure 1D-F, Supplemental Figure S5). Although carriers lacking Rab5b (and instead marked by other GTPases, e.g. Rab4a, Rab4b, Rab11) also emerge and were not visualized, the increase in Rab5b tubules is consistent with proposed roles for both IST1 and CHMP1B in scission of tubulovesicular carriers leaving the early/sorting endosome.

To evaluate additional contributions of IST1 to endosomal function, we immunostained IST1-depleted cells with an antibody that recognizes mono-and poly-ubiquitin conjugated proteins (Fujimuro *et al*., 1994) to look for abnormalities in processing cargo typically targeted into MVBs for ESCRT-dependent degradation. As expected, depleting CHMP4B increased ubiquitinated material on swollen vacuolar structures (Odorizzi *et al*., 1998; Yang *et al*., 2021) (Supplemental Figure S6A arrow). Depleting IST1, however, had no effect on the normal distribution of ubiquitin conjugates (Supplemental Figure S6B). Together with earlier studies showing that IST1 is not required for efficient EGF receptor degradation (Agromayor *et al*., 2009) this suggests that IST1 is not needed for ESCRT-dependent MVB cargo degradation.

### IST1 functions in slow but not fast transferrin recycling

Transferrin (Tfn) and its receptor (TfnR) are responsible for ongoing and essential iron uptake into cells and are widely studied because of their readily monitored trafficking through the endosomal system. Multiple pathways recycle internalized TfnR from early/sorting endosomes, classified decades ago as ‘fast’ and ‘slow’ based on kinetics, with distinct molecular players and endosomal compartments responsible for direct *vs*. indirect return to the plasma membrane (Maxfield and McGraw, 2004; Grant and Donaldson, 2009). To ask how IST1 impacts these pathways, we depleted it and examined TfnR trafficking. At steady state, immunostaining showed that TfnR was dispersed from its typical juxtanuclear localization (Figure 2A). We confirmed that this redistribution was caused by loss of IST1 using a second siRNA duplex (Supplemental Figure S7A). To monitor receptor trafficking, we followed the distribution of internalized fluorescently labelled Tfn over time (Kalaidzidis *et al*., 2015; Schindler *et al*., 2015). Strikingly, IST1-depleted cells contained only about half as much Tfn as did controls after a 1 hr incubation (Figure 2B). This decrease in steady-state Tfn accumulation was not caused by differences in total TfnR (Figure 2C), pointing instead to changes in receptor trafficking. Comparing surface exposed TfnR in control *vs*. IST1-depleted cells revealed that antibody-detected TfnR on non-permeabilized cells increased in the absence of IST1 (Figure 2D) further indicating that loss of IST1 changed receptor trafficking. Despite this difference, IST1-depleted cells internalized a similar amount of Tfn during a 30 sec pulse compared to controls (Figure 2E) and bound comparable amounts of Tfn when incubated on ice for 30 min (Supplemental Figure S7B). Consistent with the observed differences in TfnR distribution, internalized Tfn remained dispersed in IST1-depleted cells (Figure 2F; Supplemental Figure S8).

**Figure 2.**
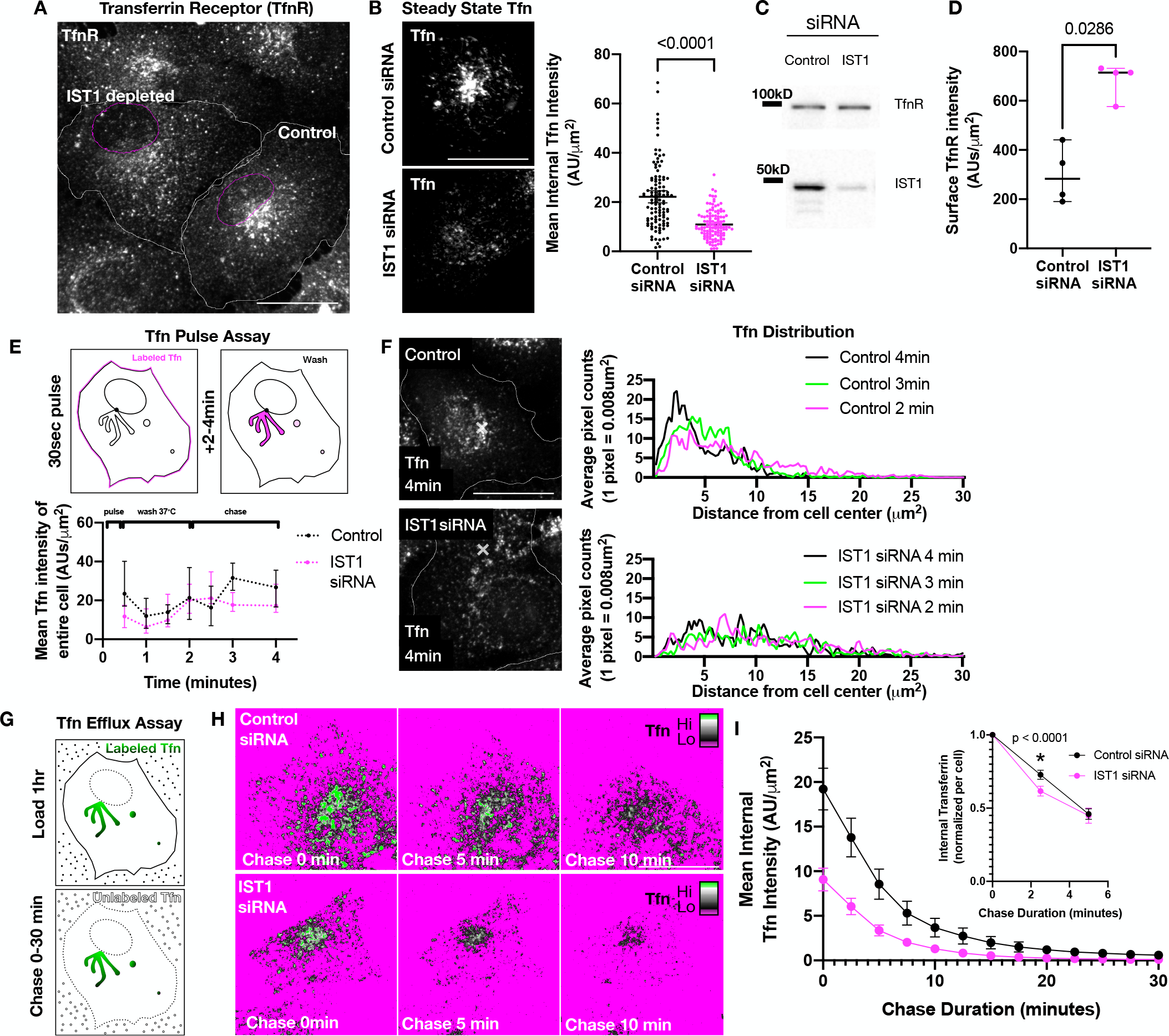
Tfn retention within the cell is impaired in IST1 depleted cells. **(A)** T24 cells individually treated with control or IST1 siRNA and then mixed and plated together and stained for TfnR (shown) and IST1 (not shown)**. (B)** T24 cells treated with control or IST1 siRNA were incubated with Tfn Alexa 555 for 1 hr, washed with PBS, and imaged immediately after adding chase media containing excess unlabeled Tfn. Quantification of mean intensity of Tfn at time 0 min. Graph shows mean± 95% CI. Control siRNA and IST1 siRNA treated cells n=110,120 (n number pooled from 3 independent experiments with n=26,25 n=42,44 n=42,51), respectively. Statistical analysis using student’s t test showed significant decrease in Tfn levels in IST1 depleted cells. **(C)** Immunoblots of TfnR and IST1 from lysates of cells depleted of IST1 or LAP1 (control) siRNA **(D)** Quantification of non-permeabilized cells immunolabeled with antibody against the extracellular domain of TfnR to assess TfnR levels on PM. Each point (n =4 for Control and IST1 siRNA treated cells) represents average TfnR intensity area for a field of cells (containing > 5 cells) normalized by the number of nuclei (i.e. average TfnR intensity per cell). Graph shows median ± 95% CI. Statistical analysis using Mann Whitney test showed significant increase in TfnR levels on PM in IST1 depleted cells. This significant increase was replicated in two more independent experiments. **(E)** Diagram of pulse Tfn assay and quantification of T24 cells allowed to internalize a 30 second pulse of Tfn 555, washed for 1.5 minutes and then chased with unlabeled Tfn. Cells were fixed at the points indicated, n=15, 12, 12, 18, 15, 21, 17 (control) and n=14, 13, 11, 17, 18, 12, 12 (IST1 KD). Graph shows median ± 95% CI. Statistical analysis using 2-way ANOVA showed Tfn uptake was not impaired in IST1 depleted cells**. (F)** Fluorescence of Tfn in IST1 siRNA treated and control T24 cells that were fixed (at the indicated times) after internalizing a 30 second pulse of Tfn 555. To compare the localization of Tfn signals, the distribution of an equivalent area (4 μm^2^) of the highest intensities was assessed by K-means clustering (see Supplemental Figure S8). The distances of each pixel from the determined cell center was plotted in a histogram. The histograms shown are an average of n=18, 21, and 17 for control and n=17,12, and 12 for IST1siRNA treated cells at 2, 3, and 4min, respectively. Similar results were observed for 3 independent experiments. All cell images are max intensity projections. **(G)** Diagram of the Tfn load-chase assay. **(H&I)** Time-dependent decay of mean Tfn intensity, measured at 2.5 min intervals after the start of the chase**. (**H) shows selected time points of 0,5, and 10 minutes and (I) is quantification of all timepoints, n=26 for control siRNA and n=25 IST1 siRNA cells. Error bars are SEM. Inset in I is internal Tfn Intensity normalized for each cell for 0, 2.5, and 5 min timepoints. Control siRNA and IST1 siRNA treated cells n=110,120 (3 independent experiments with n=26,25 n=42,44 n=42,51), respectively. **Graph shows mean** ± 95% CI. Statistical analysis using 2 way ANOVA showed Tfn exocytosis was faster at 2.5 minutes in IST1 depleted cells. Scale bars in whole cells in (A), (B), (F) and (H) are 25 µm.

Changes in the proportion of peripheral *vs*. juxtanuclear Tfn and TfnR have previously been observed when proteins important for slow recycling (e.g. indirect *via* recycling endosomes) including SNX4 (Traer *et al*., 2007), aftiphilin (Hirst *et al*., 2005), Rabenosyn-5 and EHD-1 (Naslavsky *et al*., 2004) are depleted. This suggested that IST1 might also preferentially contribute to slow recycling. Indeed, earlier studies showing elongated SNX1-and SNX4-marked tubules in IST1-depleted cells (Allison *et al*., 2013) support a role for IST1 in creating and/or releasing carriers that traffic from sorting to recycling endosomes. Decreases in slow recycling are typically offset by compensatory increases in direct or rapid recycling to the plasma membrane, so we next asked if depleting IST1 changed the rate of Tfn efflux. To monitor this, we chased cells loaded with fluorescent Tfn with unlabeled Tfn and quantified fluorescence remaining as a function of time (Figure 2G-I). IST1-depleted cells again accumulated only half as much Tfn as did controls. Comparing changes in normalized Tfn intensity of individual cells over time revealed accelerated early loss of Tfn in cells depleted of IST1 (Figure 2I, inset). Altogether these results point to a role for IST1 in slow TfnR recycling.

A limitation to the sensitivity of our Tfn recycling assays arises from the different affinities of holo-Tfn (iron bound) and apo-Tfn (iron free) for TfnR (Yersin *et al*., 2008), since changes in TfnR trafficking within the endosomal system (i.e. reaching a sufficiently acidic endosome for iron release from Tfn) may affect iron assimilation (Chen *et al*., 2013) and thus the subsequent exchange of unlabeled for labeled Tfn. This could explain why IST1 depleted cells have more TfnR on their surface (Figure 2D) but bind and internalize the same amount of labeled Tfn as controls (Figure 2E, Supplemental Figure S7B). Newly developed methods to rapidly detect TfnR itself, such as using HaloTag ligands (Jonker *et al*., 2020), will be a way to further study the many factors that together control TfnR recycling.

### IST1-independent recycling involves endosomes enriched in AP-1

Recycling pathways leaving the early/sorting endosome depend on many factors for cargo sorting and carrier biogenesis. Prominent among these are retromer, retriever, COMMD/CCDC22/CCDC93 (CCC), endosomal SNX-BAR sorting complex for promoting exit-1 (ESCPE-1) and the actin nucleation promoting Wiscott–Aldrich syndrome protein and SCAR homologue (WASH) complex. Clathrin and associated machinery also contribute. IST1 is not known to directly bind any of these factors. To better understand the role(s) of IST1 in slow *vs*. fast TfnR recycling we set out to differentiate pathway(s) that are or are not affected by depleting IST1. Given earlier reports of impaired SNX1-and SNX4-marked carrier scission (Allison *et al*., 2013, 2017) and the defect in slow TfnR recycling described above, we focused on clathrin and its adaptor AP-1 that have long been known to participate in rapid recycling (Hirst *et al*., 2005, 2012; D’Souza *et al*., 2014; Chamberland *et al*., 2016). After depleting IST1 we observed a striking redistribution of AP-1 from its typical perinuclear concentration to the cell periphery (Figure 3A). To ask if TfnR accesses these AP-1 marked compartments, we compared Tfn localization during initial uptake to that of AP-1 γ-adaptin. After 2-3 minutes, Tfn in IST1-depleted but not control cells remained in peripherally distributed endosomes which were robustly stained for AP-1 (Figure 3A-B). Coincident AP-1 and Tfn were apparent on both punctate and tubular organelles (Figure 3B), suggesting that without IST1 TfnR recycles *via* a pathway similar to the clathrin-coated vesicle lined tubular-vesicular networks described many years ago by whole-mount electron microscopy (Stoorvogel *et al*., 1996).

**Figure 3.**
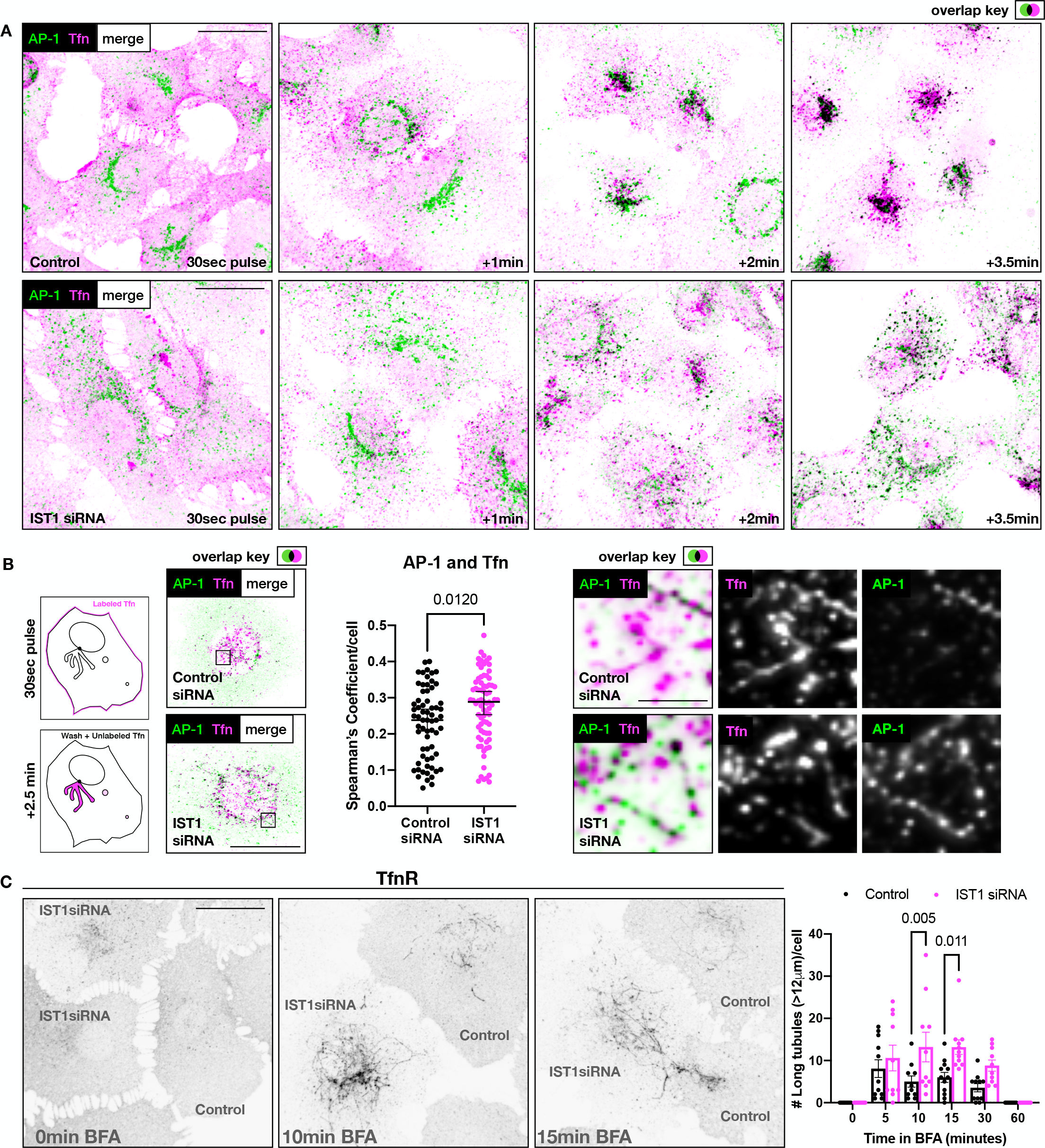
Dispersed Tfn positive compartments in IST1 depleted cells are associated with the endosomal clathrin adaptor AP-1. **(A)** Control (upper panel) or IST1 depleted T24 cells (lower panel) were treated with 30 second pulse of Tfn555, washed and allowed to continue endocytosis of bound Tfn for 1.5 minutes before being chased with unlabeled Tfn. Cells were fixed at 30 seconds (1st panel), 1.5 minutes (2nd panel), 2.5 minutes (3rd panel) or 4 minutes (4th panel) and stained for AP-1. **(B)** Diagram of a short Tfn pulse, where a 30 sec pulse of labeled Tfn is immediately followed by chase media, and examples of Control KD and IST1 KD cells where IST1 KD cells show AP-1 localization along abnormal peripheral Tfn tubules. The Spearman’s Correlation Coefficient between AP-1 and Tfn was quantified for three independent experiments, total of n= 65 and n= 76 for control and IST1 siRNA treated cells. Graph shows median and 95% ± CI, and statistical analysis using Mann-Whitney test showed significant increase in overlap of AP-1 and Tfn in IST1 depleted cells **(C)** Mixed IST1 siRNA and control T24 cells were fixed before and after treatment with 10 µM brefeldin A (BFA) for 0, 10, 15 minutes and stained for TfnR (shown) and IST1 (not shown). All cell images are max intensity projections. Tubules longer than 12 μm within tubular networks were quantified at 0,5,10,15,30,60 minutes. n ζ 10 for each condition, Statistical analysis using 2-way ANOVA demonstrated increased tubulation in IST1 depleted cells at 10 and 15 min. Scale bars in whole cells in (A), (B) and (C) are 25 µm, scale bar in insets in (B) is 2.5 μm.

To further characterize IST1-independent rapid recycling, we took advantage of the selective effects of brefeldin A (BFA) on different elements of the endosomal system. BFA releases Arf1-activated AP-1 from membranes (Robinson and Kreis, 1992; Wong and Brodsky, 1992), tubulating Rab4a-containing endosomes involved in rapid recycling with little or no effect on the Rab11 endosomes associated with slow recycling (Daro *et al*., 1996; Stoorvogel *et al*., 1996). We found that treating T24 cells with BFA transiently redistributed TfnR from the cell surface into extended tubular compartments (Figure 3C) highlighting a role for BFA-sensitive pathway(s) in TfnR recycling. Acutely treating IST1-depleted cells with BFA led to even more extensive tubular networks that took longer than their control counterparts to resolve (Figure 3C), consistent with an enhanced need for AP-1 dependent carriers and the rapid recycling pathway they delineate.

For further insight into how IST1 affects cargo exit from early/sorting endosomes, we asked how depleting it affects mannose 6-phosphate receptor (M6PR) trafficking as this receptor has also been reported to move through both retromer and AP-1 marked carriers (Arighi *et al*., 2004; Saint-Pol *et al*., 2004; Seaman, 2004; Popoff *et al*., 2007, 2009). M6PR’s primary itinerary involves anterograde export from TGN to endosomes and retrograde transit from various endosomes back to the TGN. At steady state, M6PR is typically concentrated in the perinuclear TGN. Allison *et al*. have previously reported that loss of IST1 impairs return of M6PR to the TGN (Allison *et al*., 2017). We found that cells depleted of IST1 often contained dispersed M6PR some of which overlapped with EEA1 (Figure 4). Using super-resolution Airyscan confocal imaging, we examined the distribution of M6PR and EEA1 in more detail (Figure 4B). We found that while tubules containing M6PR originated from EEA1 endosomes in both IST1 depleted and control cells, overlap of M6PR and EEA1 on the endosomal perimeter was increased in cells depleted of IST1. This suggests that IST1 helps to efficiently concentrate M6PR in tubules that leave the sorting endosome. Altogether, the changes in TfnR and M6PR trafficking caused by depleting IST1 point to an important role for this ESCRT-III protein on the early/sorting endosome in events needed to create and/or release carriers destined for recycling endosomes and/or the TGN. Future studies will be needed to examine the effect of blocking IST1 on the structure and organization of recycling endosomes. The shift from slow to rapid recycling seen in IST1-depleted cells highlights the plasticity of trafficking events on sorting endosomes.

**Figure 4.**
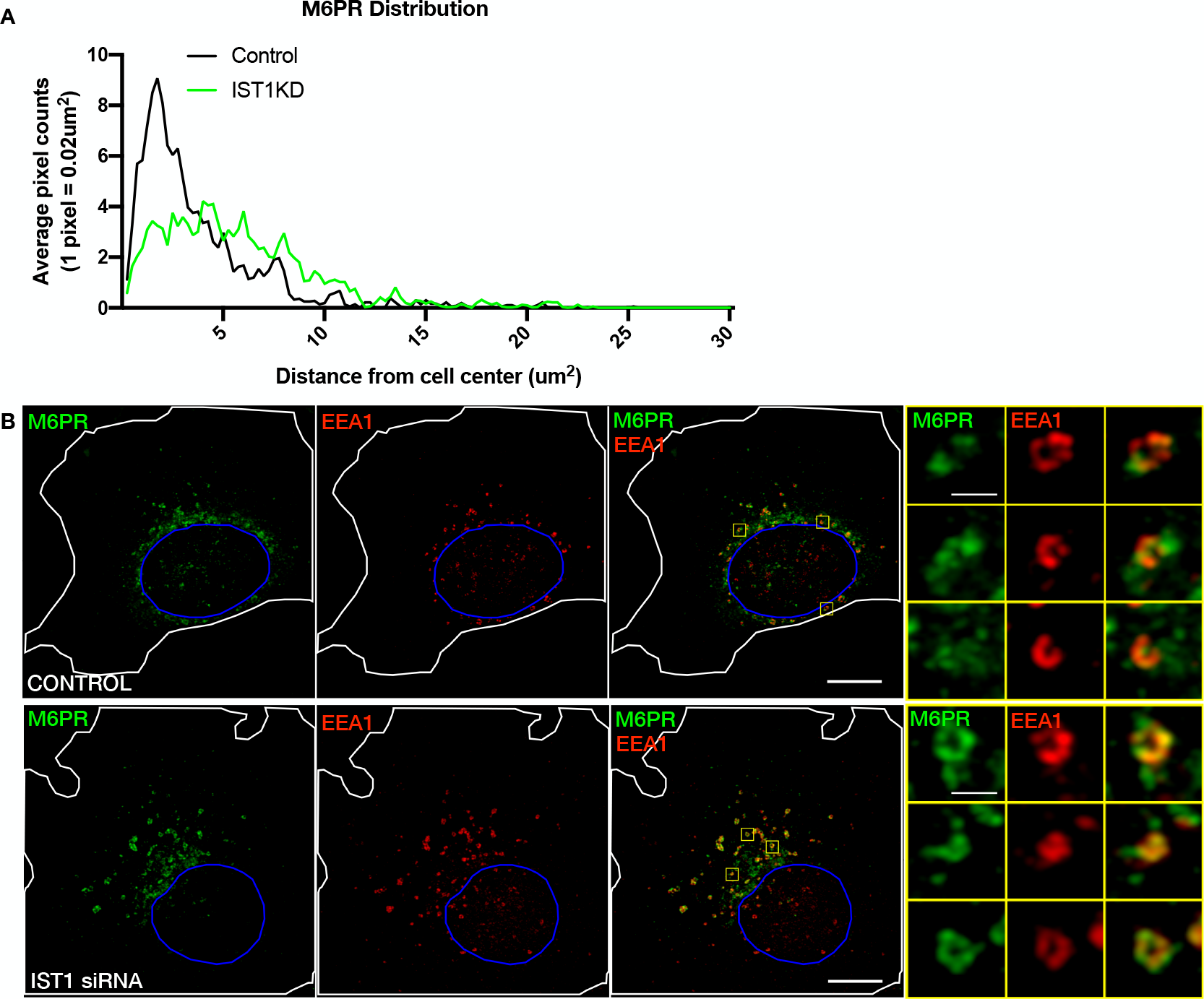
IST1 depletion causes M6PR to accumulate in EEA1-labeled early/sorting endosomes. **(A)** To compare the localization of M6PR signals, the distribution of an equivalent area (4 μm^2^) of the highest intensities was assessed by K-means clustering (as shown previously for Tfn in Supplemental Figure S8). The distances of each pixel from the determined cell center was plotted in a histogram. The histograms shown are an average of n=25 for control and IST1siRNA treated cells (data shown is compiled from 3 independent experiments with each demonstrating increased dispersed M6PR in IST1 siRNA treated cells). (**B**) Immunostaining of M6PR and EEA1 in control and IST1 depleted cells. Images are maximum projections of z-stacks obtained on Zeiss LSM 880 with Airyscan and subsequent Airyscan processing. Contrast adjustments on M6PR channel are equivalent for both conditions, but are different for the EEA1 channel, due to increased EEA1 recruitment in IST1 depleted cells. Scale bars in the whole cells and insets are as follows: (B) 10 μm and 1 μm, respectively.

### IST1 and SNX15 occupy a distinct subdomain on sorting endosomes

Given the role of IST1 in a subset of pathways leaving early/sorting endosomes we next wanted to know when and where it is present on these organelles. To kinetically define them, we incubated cells with fluorescently labelled Tfn for 2-3 minutes. Immunostaining for EEA1 and IST1 revealed that newly internalized Tfn was present in compartments frequently marked by both of these proteins (Figure 5A). Notably, discrete spots of IST1 were adjacent to but not overlapping with EEA1. Once endosomes acquire EEA1, confocal fluorescence microscopy can usually resolve “degradative” and “retrieval” subdomains defined respectively by HRS (ESCRT-0) or retromer and associated sorting nexins together with branched actin (Cullen and Steinberg, 2018). To ask whether IST1 coincides with markers of either of these subdomains we compared its localization with that of HRS and the retromer protein VPS35. We also examined sorting nexin 15 (SNX15), a protein previously implicated in regulating recycling (Feng *et al*., 2016) and present on newly formed endosomes (Danson *et al*., 2013; Flores-Rodriguez *et al*., 2015). Immunostaining revealed that IST1 did not colocalize well with HRS or VPS35 but appeared to overlap with SNX15 (Figure 5B). However, the juxtanuclear concentration of endosomes made it difficult to examine the relationships among these factors in detail.

**Figure 5.**
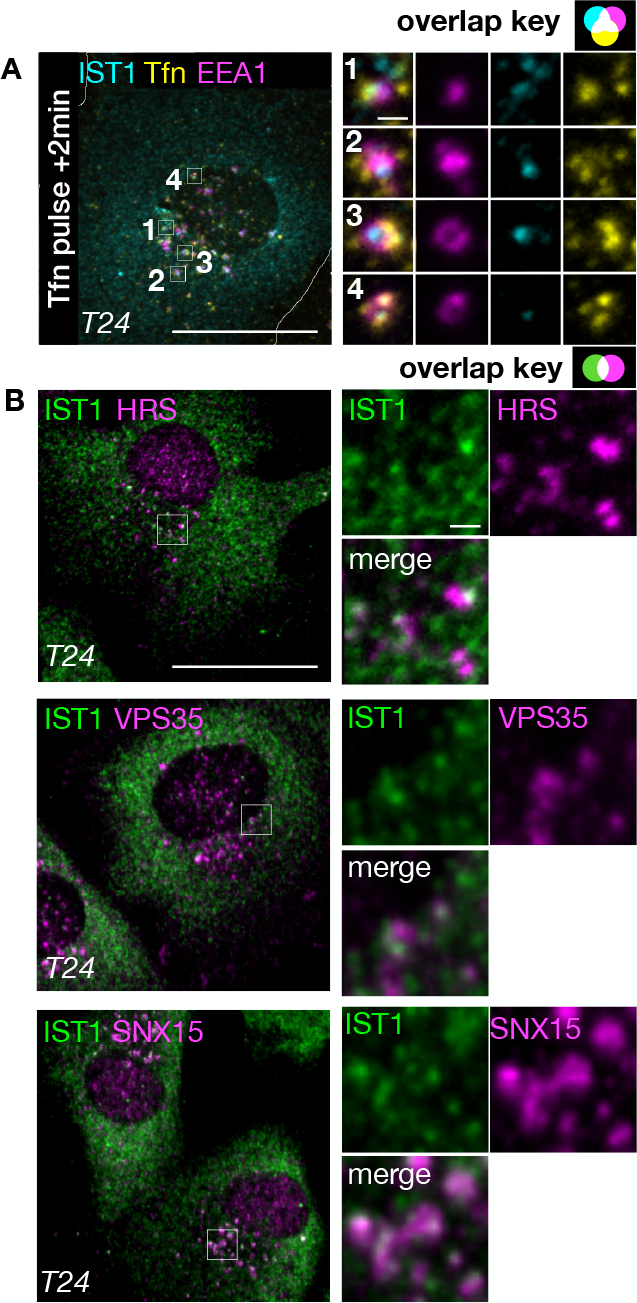
IST1 is proximal to early endosome markers HRS, VPS35, and SNX15. IST1 is proximal to recently endocytosed Tfn and known early/sorting endosome domain markers. **(A)** T24 cells allowed to internalize a 30 second pulse of Tfn 555 and were fixed after 2.5 min and stained for IST1 and EEA1. **(B)** T24 cells stained for IST1 and either HRS, VPS5 or SNX15. All cell images are max intensity projections. Whole cells and insets are as follows: (A) 25 μm and 1 μm, (B) 25 μm and 1 μm, respectively.

To spatially and temporally resolve endogenous proteins as they appear on nascent endosomes, we depolymerized microtubules to slow endosomal movement and block perinuclear clustering. We then monitored the presence of proteins on individual endosomes marked by newly internalized cargo (Derivery *et al*., 2009, 2012). As an endocytic tracer, we used fluorescently labelled wheat germ agglutinin (WGA), a lectin which binds to sialic acid and N-acetylglucosamine (GlcNAc) moieties on plasma membrane proteins (Chazotte, 2011) (Raub *et al*., 1990). We internalized pre-bound WGA for 5 min, fixed and immunostained cells, and then examined the relationship among different signals on endosomes (regions of interest (ROIs)) identified by internalized WGA (Figure 6). Most, but not all, WGA containing ROIs were marked by EEA1 (Figure 6C empty arrow head, Supplemental Figure S9A), and approximately 60% of EEA1-marked ROIs acquired WGA during the 5 min internalization window (Figure 6C filled arrow heads, Supplemental Figure S9A).

**Figure 6.**
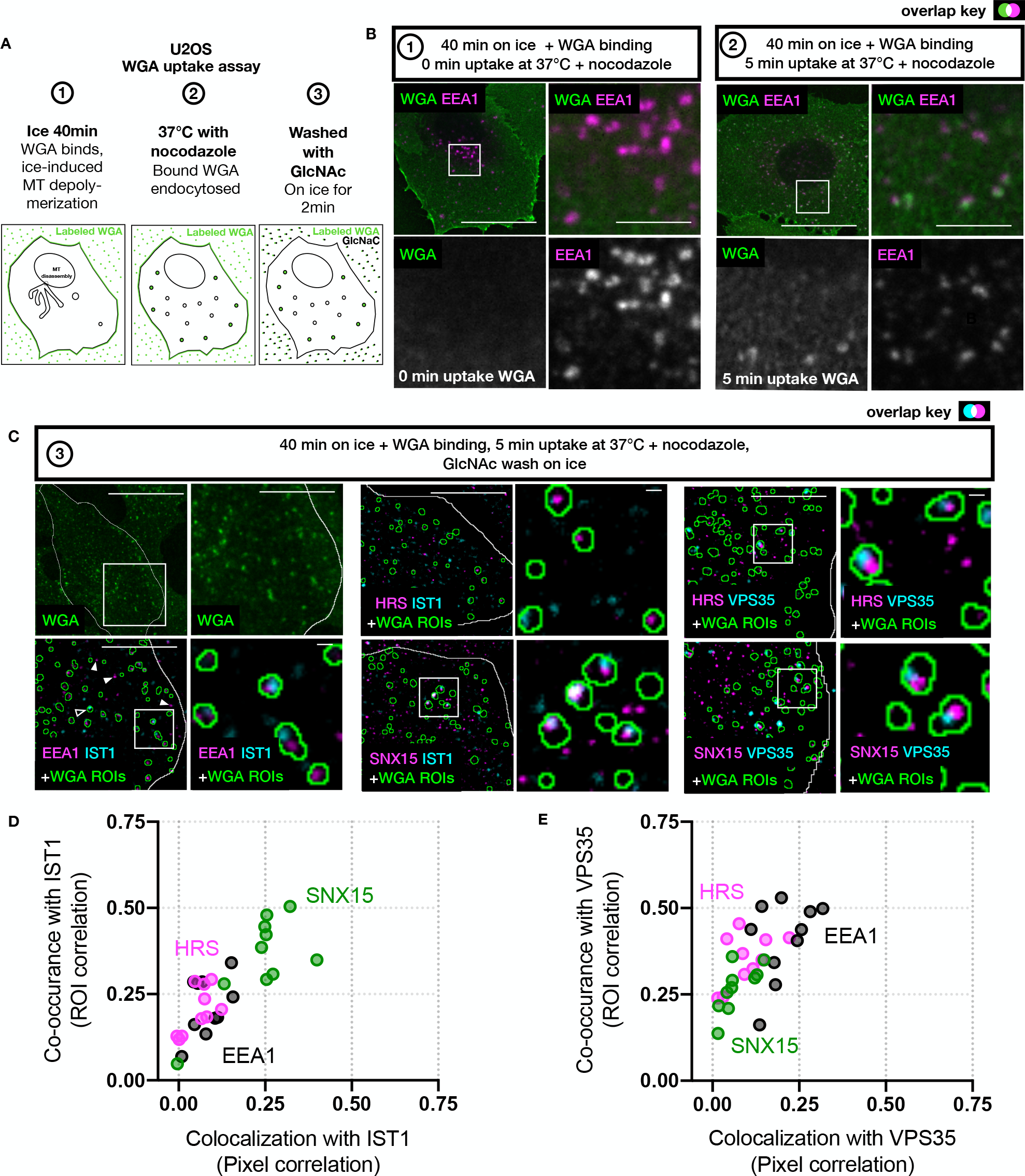
IST1 distribution to subdomains on early/sorting endosomes. **(A)** Schematic of the WGA assay. U2OS cells are incubated on ice with WGA-TRITC for 30 min and either fixed in PFA **(B, left**), or (2) allowed to internalize bound WGA for 5 min at 37 °C in the presence of nocodazole **(B, right),** and then stained for EEA1. Images in B are single slices. (**C**) Cells treated identically as B, but (3) washed in ice cold GlcNAc prior to fixation to remove remaining plasma membrane associated WGA, and then stained for two makers. The first set of panels shows WGA signal, which was used to generate WGA ROIs (green outlines) shown with EEA1 and IST1 at the bottom. The remaining two sets of panels show either IST1 or VPS35 co-stained with either HRS or SNX15. Cell images are max intensity projections. (**D**) A plot of the Spearman’s rank coefficient of pixel-based vs ROI-based correlations of IST1 relative to EEA1 (n=10), HRS (n=12) or SNX15 (n=10). **(E)** A plot of the Spearman’s rank coefficient of pixel-based vs ROI-based correlations of VPS35 relative to EEA1 (n=10), HRS (n=10) or SNX15 (n=10). Scale bars in the whole cells and/or insets are as follows: (B) 25 μm and 5 μm, respectively; (C) 25 μm, 5 μm, and 1 μm respectively.

Using this experimental paradigm we interrogated the relationship between IST1 and markers of endosomal subdomains. A commonly used approach for assessing correlation is to determine if there is a linear relationship between the intensity of two signals across individual pixels in an image (i.e. Spearman’s rank coefficient or Pearson’s correlation coefficient). This analysis will, however, miss relationships between signals that are present or juxtaposed within a ROI but not directly overlapping. These can be described using ROI-based analysis of the correlation between two signals present within the same endosome or ROI. By comparing pixel-and ROI-based correlations we can distinguish co-localization vs. co-occurrence of relevant proteins within endosomal ROIs. With this approach, we found that IST1 had poor pixel-based correlation (i.e. co-localization) with EEA1 or HRS (Figure 6D, x axis) and higher pixel-based correlation with SNX15. Notably, IST1 was better correlated with EEA1 and HRS in ROI-based analyses (Figure 6D, y axis; Supplemental Figure S9B), indicating that these proteins are juxtaposed but not overlapping.

We next explored the relationship between SNX15 and factors previously implicated in cargo retrieval/recycling (Cullen and Steinberg, 2018). We examined VPS35 relative to HRS, SNX15 and EEA1. VPS35 showed poor pixel-based correlation (Figure 6E, x axis) and increased ROI-based correlation (Figure 6E, y axis) with all three signals. This confirmed earlier reports of a proximal but non-overlapping relationship between HRS and VPS35 on early/sorting endosomes (Norris *et al*., 2017; MacDonald *et al*., 2018). Additionally the juxtaposed relationship we find between VPS35 and SNX15 is similar to what has previously been reported for IST1 and SNX1 (Allison *et al*., 2017). Taken together, there appear to be at least three separable but adjacent subdomains on early/sorting endosomes – one marked by HRS, one by VPS35, and one by SNX15 and IST1.

### IST1 co-assembles with SNX15 on endosomes

Given the presence of IST1 and SNX15 in a common subdomain on the early/sorting endosome, we next asked whether they directly interact. IST1, like other ESCRT-III proteins, has C-terminal “MIM” motifs that bind to MIT domains to mediate specific protein-protein interaction. SNX15 is a MIT domain-containing protein recruited to endosomes via a PX domain that binds phosphatidyl-inositol-3-phosphate (Phillips *et al*., 2001; Danson *et al*., 2013) (Figure 7B). Its MIT domain shares ∼30% homology with MIT domains in VPS4A/VPS4B but had until a recent *in vitro* study of the human ESCRT-III-MIT domain “interactome” (Wenzel *et al*., 2022) not been shown to bind ESCRT-III proteins. To explore a possible interaction between SNX15 and IST1 in cells, we co-overexpressed them in U2OS cells. Consistent with earlier yeast two-hybrid interaction analyses (Danson *et al*., 2013), co-overexpressed ESCRT-III proteins CHMP1 – CHMP7 did not co-localize with SNX15 (data not shown). IST1-mApple, however, strikingly accumulated with SNX15-GFP on discrete structures spread across the cell (Figure 7A, inset Figure 7A). The presence of internalized Tfn within these structures, as well as within protruding tubules, confirmed their identity as endosomes (Figure 7A).

**Figure 7.**
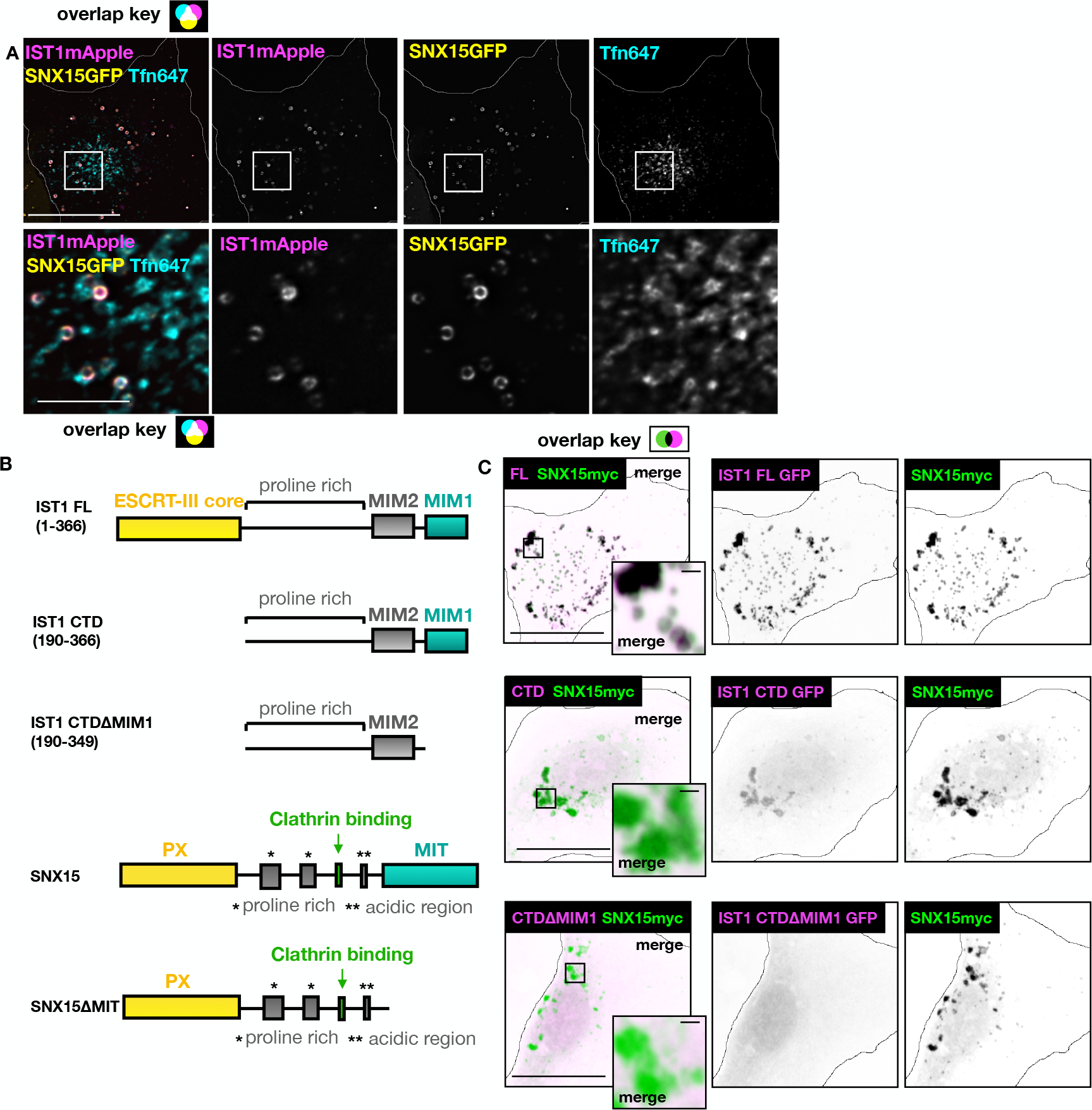
IST1 uniquely associates with SNX15. **(A)** IST1 and SNX15 co-assemble on Tfn-containing endosomes in living U2OS cells. Cell images are single slices. **(B)** Domain diagram of IST1 (top), SNX15 (bottom), and constructs used to examine the interaction between IST1 and SNX15. **(C)** Fixed cells expressing SNX15myc with GFP-tagged IST1 constructs shown in B, in the inverted magenta-green images overlapping signal is black and captures dynamic range of IST1 constructs recruited to SNX15. Cell images are max intensity projections. Scale bars in the whole cells and insets are as follows: (A) 25 μm and 5 μm, respectively; **(A)** (C) 25 μm and 1 μm, respectively.

If SNX15 helps bring IST1 to the membrane, we predicted that its overexpression would also recruit endogenous IST1 to the endosome. Indeed, overexpressing SNX15 redistributed IST1 onto endosomes while depleting it from the cytoplasm (Supplemental Figure S10A). To determine how IST1 engages SNX15, we asked if removing its N-terminal “core” ESCRT-III domain affected assembly with SNX15 (Figure 7B). We found that a C-terminal fragment (CTD) of IST1 was still recruited to endosomes by over-expressed SNX15, albeit to a lesser extent than full-length IST1 (Figure 7C). To further delineate sequences responsible for this interaction, we deleted the C-terminal MIM1 motif from IST1-CTD and found that this abolished its SNX15-mediated recruitment to endosomes (Figure 7C). Conversely, we removed the MIT domain from SNX15 and found that this fragment no longer depleted IST1-mApple from the cytoplasm (Supplemental Figure S10B). Together with the recently reported biochemical characterization of binding between SNX15’s MIT domain and IST1 (Wenzel *et al*., 2022), these observations indicate that interaction between IST1’s MIM1 and SNX15’s MIT domains provides a novel mechanism by which IST1 can be recruited to early/sorting endosomes independently of other ESCRT proteins.

### SNX15 and CHMP1B alternately distribute IST1 between endosomal subdomains

Having found that SNX15 and IST1 co-assemble on early/sorting endosomes around much of the endosomal perimeter, we next examined their relationship with other endosome-associated proteins (Figure 8A-D). Using cells stably expressing low levels of SNX15GFP, we found that similar to the relationship between endogenous SNX15 and VPS35 (Figure 6), other proteins involved in retrieval such as SNX1, the WASH complex (FAM21 and WASH), and branched actin-associated cortactin colocalized poorly with SNX15GFP as assessed by Pearson’s correlation coefficient but were nonetheless present on the majority of SNX15GFP containing endosomes (Figure 8E, insets in Figure 8A-D). Overexpressing SNX15 thus expands its coverage of the endosomal perimeter but preserves the subdomain relationships that we observed with endogenous proteins on dispersed endosomes (Figure 6).

**Figure 8.**
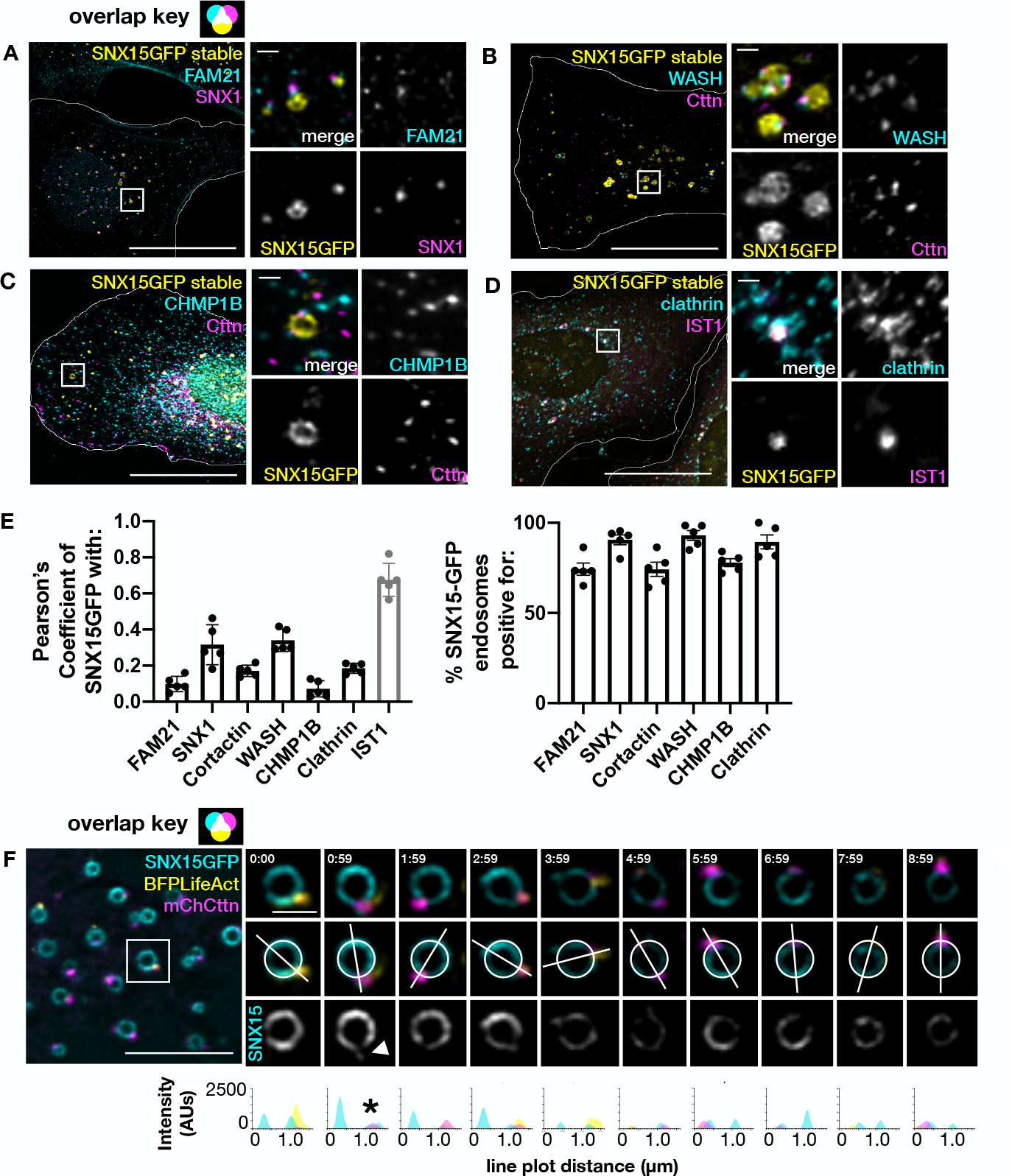
IST1 assembled with CHMP1B localizes to tubules associated with endosomes, which is distinct from the localization of SNX15. **(A-D)** SNX15gfp stable cells were fixed either with methanol +BS3 (A, C and D) or with PFA (B), and stained with the indicated antibodies of FAM21/SNX1, WASH/cortactin, CHMP1B/cortactin, or IST1/clathrin. All images were deconvolved using Nikon software. Cell images (A-D) are max intensity projections. (E) Quantification of the pearson’s correlation coefficient of SNX15GFP and the indicated marker (left) and manual quantification of the presence of the indicated marker (except for IST1) proximal to SNX15gfp endosomes defined as ROIs (right). (F) Live cell imaging, deconvolved using Nikon software. U2OS cell expressing exogenous SNX15gfp, BFP LifeAct, and mcherry cortactin. Panel to the left shows time-lapse imaging of inset SNX15GFP endosome, as well as line scans that bisect the endosome and the actin domain. Images are single slices. Scale bars in whole cells and insets are as follows: (A-D) 25 μm and 1 μm (F) 5μm and 1 μm, respectively.

Because both SNX15 and CHMP1B bind directly to endosomal membranes (Phillips *et al*., 2001; McCullough *et al*., 2015), we next wondered if they cooperate to engage IST1. This seemed possible because of their distinct interactions with IST1, with CHMP1B binding to the core ESCRT-III domain (Bajorek *et al*., 2009b) and SNX15 its C-terminal MIM motifs (see above). Interestingly, we found that there was no CHMP1B apparent within SNX15 enriched domains (Figure 8A), despite the abundant recruitment of IST1 (Figure 8D). This suggests an alternative model in which IST1 assembles with CHMP1B only at sites devoid of SNX15, potentially where tubules emerge from endosomes. To determine if this is the case, we compared the distribution of CHMP1B to that of markers of branched actin that localizes to the base of endosomal tubules (Cullen and Steinberg, 2018). Indeed, we saw examples of CHMP1B coincident with cortactin (Figure 8C). Using live cell imaging, we confirmed that over time SNX15GFP remained concentrated on the vacuolar endosome, with an intensity that decreased near and around puncta marked by actin (Figure 8F) as well as in transient “fingers” of signal extending beyond the actin puncta onto connected tubules (Figure 8F, arrow and asterisk). Altogether, these results indicate that IST1 bound to SNX15 resides primarily on the endosomal perimeter and does not overlap with CHMP1B.

Within the endosomal pathway, clathrin contributes to recycling but also forms a “flat” clathrin-containing domain on early/sorting endosomes that regulates the dynamics of ESCRT-0/HRS and ILV formation (Wenzel *et al*., 2018). SNX15 has been reported to bind clathrin (Danson *et al*., 2013) and we wondered about its relationship (alone or together with IST1) with endosome associated clathrin. Although the overall correlation between SNX15GFP and clathrin in cells was relatively low (Figure 8E), they clearly overlapped on vacuolar endosomes and connected tubules (Figure 8D&E). To explore the dynamics of this relationship and potential roles for SNX15 in regulating endosomal clathrin, we examined live cells transiently expressing mRFP clathrin light chain A (CLCa) and SNX15GFP and confirmed some overlap (Figure 9A arrow) with a higher Pearson’s correlation coefficient than seen with endogenous proteins (Supplemental Figure S11A&B). Clathrin was present with SNX15 on the endosomal perimeter (Figure 9A arrow) as well as on tubular and punctate structures originating from the same endosomes (Figure 9A arrowheads). Tfn in the clathrin-marked structures (Supplemental Figure S11C) confirmed their endosomal origin.

**Figure 9.**
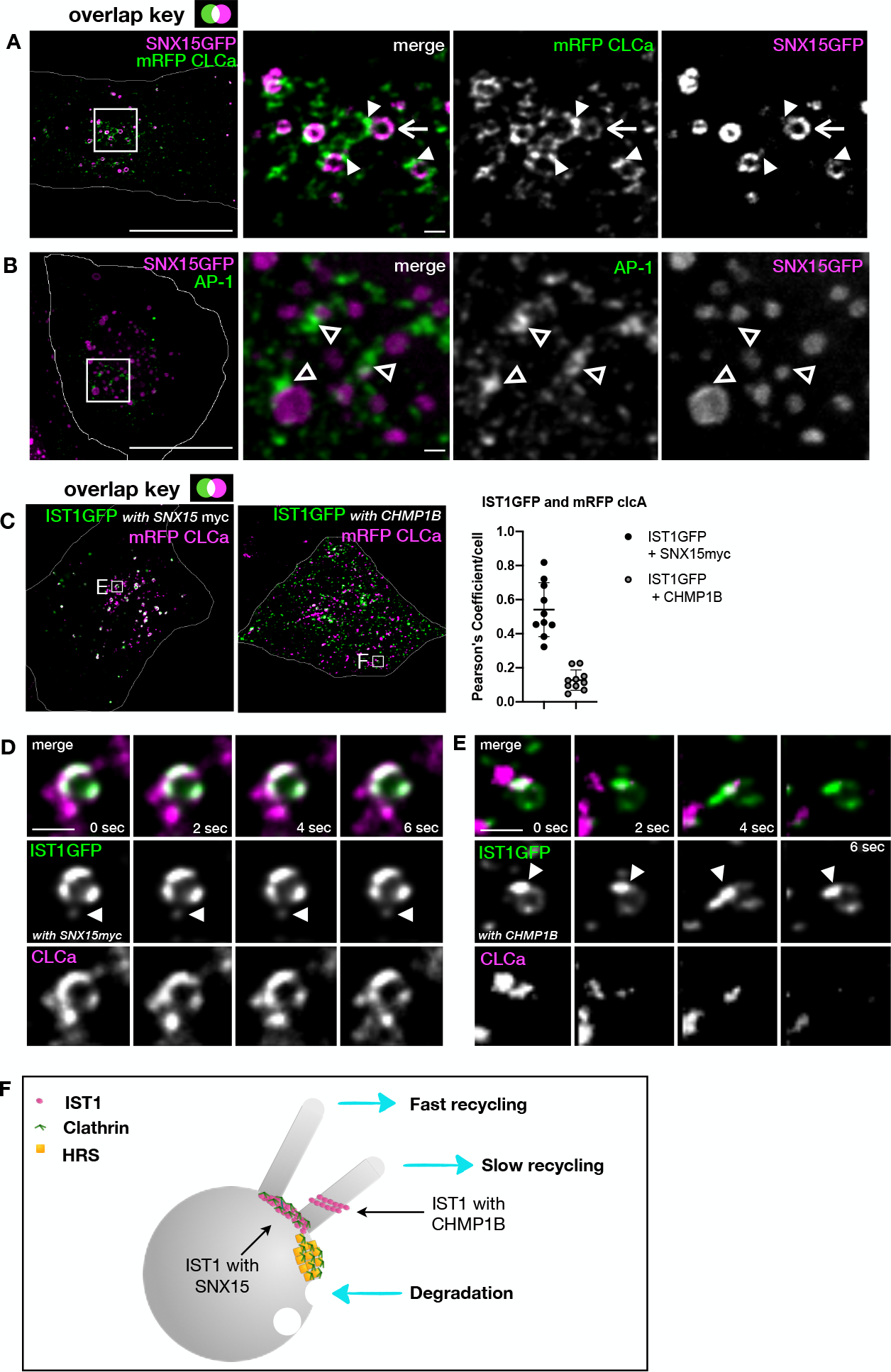
IST1 functions on early/sorting endosomes in tubule scission. **(A)** U2OS cell stained for endogenous IST1, asterisk denote IST1 at cytokinetic cell bridges and mid-bodies. **(B)** U2OS (right) and T24 (left) cells immunostained with IST1 and EEA1. **(C)** Immunostaining of EEA1 in control and IST1 siRNA treated T24 cells, and the recruitment of EEA1 is quantitated to the right, n= 17 for control, n=19 for IST1siRNA treated. Graph shows median ± 95% CI. Statistical analysis with Mann Whitney U test showed significant increase in EEA1 in IST1 depleted cells. These results were observed for 3 independent experiments. Images in (A-C) are max intensity projections. **(D-F)** Stable U2OS cell lines expressing GFP-Rab5 were treated with IST1, CHMP1B, or the unrelated protein LAP1 (control) siRNA and Rab5 tubule number and duration were assessed by live cell imaging. **(D)** Representative single frames, single slices from live cell movies. Note the presence of long tubules (inset) in cells depleted of CHMP1B and IST1. **(E)** Each dot in the scatter plots shows the number of tubules in a single cell longer than 5 µm and present for >2 sec over a 30 sec recording, n=50 for all three groups (results are pooled from three independent experiments). Graph shows median ± 95% CI. Statistical analysis using Kruskal-Wallis test revealed increased number of tubules per cell for IST1 and CHMP1B depleted cells. **(F)** Bar graph totals tubules counted in (E) and further shows how long each was present (tubule duration). Not only did CHMP1B and IST1 KD cells have more tubules, but only 5 tubules observed in control KD cells persisted for over 4 seconds while 43 and 14 tubules observed in CHMP1B and IST1 KD cells, respectively, persisted for over 4 seconds. Scale bars in the whole cells and insets are as follows: (A) 25 µm, (B) 25 µm and 1 µm, (C) 25 µm, and (D) 25µm and 5 µm, respectively.

Clathrin that engages with ESCRT-0/HRS is not associated with AP-1 or other cargo adaptors (Raiborg *et al*., 2001). SNX15-coincident clathrin also did not overlap with AP-1, which was instead concentrated in adjacent, likely contiguous, structures (Figure 9B). SNX15 similarly did not colocalize with other clathrin adaptors (GGA2, GGA1, EpsinR) (data not shown), suggesting that SNX15-associated clathrin also does not form cargo carriers. However, determining whether SNX15-associated clathrin forms a “flat” domain comparable to that associated with HRS or instead a contiguous assembly of clathrin coated buds that lack the adaptors we examined will require future study using electron microscopy.

Given that IST1 function in TfnR trafficking from early/sorting endosomes appears to be compensated by separable from processes that require clathrin, we next queried the relationship between IST1 and clathrin when it was assembled with SNX15 or CHMP1B. We examined the relationship between IST1GFP and mRFP CLCa in the presence of co-overexpressed SNX15 or CHMP1B (Figure 9C-E). While IST1GFP assembled with SNX15 mainly co-localized with clathrin on the vacuolar domain, it was also present in dim “finger” structures that coincided with bright CLCa signal (Figure 9D). In contrast, IST1GFP coexpressed with CHMP1B localized predominantly to tubular structures protruding from endosomes that were transiently enriched in clathrin (Figure 9E). Moreover, although both complexes of ISTGFP overlapped with CLCa the colocalization was much greater when IST1 was assembled with SNX15 (Figure 9C). Altogether, live-cell imaging of IST1GFP in different complexes indicates that IST1 has multiple functions including a role in the dynamics of a SNX15-clathrin domain with potential contributions to sorting and a separate role in CHMP1B mediated tubule scission. Our results suggest that IST1 is alternately recruited to different subdomains of the endosome and plays a particularly important role in the carrier biogenesis needed to deliver cargo into the slow recycling pathway (Figure 9F).

## Discussion

Functional studies in both mammalian cells and yeast have previously uncovered roles for IST1 in endosomal recycling (Allison *et al*., 2013; Allison *et al*., 2017: MacDonald and Piper, 2017; Laidlaw *et al*., 2022). While initially unexpected because of the well-established roles for ESCRT proteins in MVB biogenesis and delivery to the lysosome, it is now clear that ESCRTs contribute to a much broader range of membrane remodeling activities. The ability of IST1 to polymerize around and constrict tubules coated with CHMP1B (McCullough *et al*., 2015) suggested it could act by promoting scission of endosomal tubules, either on its own or by enabling friction driven scission as recently described *in vitro* (Cada et al., 2022). Where and how IST1 cooperates with other machinery involved in endosomal sorting and scission has been explored in various settings but because of the complexity and varying specializations of the endosomal system in different cells and organisms this remains to be clearly understood, leading to the studies reported here.

Our analysis of Rab5 tubule dynamics confirmed previously described involvement of IST1 and CHMP1B in endosomal tubule scission, similar to the WASH complex (Derivery *et al*., 2009) and potentially controlled by contact with the endoplasmic reticulum (Rowland *et al*., 2014) or microtubules (Cada *et al*., 2022). Using TfnR trafficking to assess specific functional roles, we found that depleting IST1 selectively impaired TfnR movement from early/sorting endosomes into the “slow” or indirect recycling pathway. This was accompanied by increased flux through a “fast” pathway that directly recycled receptor to the plasma membrane. Notably, IST1-independent “fast” recycling involved peripheral endosomes marked by the clathrin adaptor AP-1, which is known for its contribution to fast recycling (Hirst *et al*., 2005, 2012; D’Souza *et al*., 2014; Chamberland *et al*., 2016). Indeed, depleting either the γ subunit of AP-1 (Hirst *et al*., 2005) or the AP-1 accessory protein NECAP-2 (Chamberland *et al*., 2016) impairs rapid TfnR recycling. IST1 is not the first protein selectively implicated in slow TfnR recycling; others with similar roles include SNX4 (Traer *et al*., 2007), aftiphilin and p200/HEAT repeat-containing protein 5b (HTR5B) (Hirst *et al*., 2005) and EHD1/3 (Naslavsky *et al*., 2004). We also found that M6PR, a cargo that recycles between endosomes and the TGN, accumulated at the perimeter of EEA1-marked compartments, again consistent with a need for IST1 in releasing specific cargo and cargo carriers from early/sorting endosomes, as previously shown for SNX1-marked tubules (Allison *et al*., 2013, 2017). Interestingly Laidlaw *et al*. recently reported that Ist1p and Vps4p in *Saccharomyces cerevisiae* function on a recycling endosome from which nutrient transporters return to the cell surface (Laidlaw *et al*., 2022). Its activity in this setting is regulated by post-translational modifications that have not been evident in studies of mammalian IST1 but have also not been systematically explored. Overall, IST1 seems likely to be involved in multiple events that share a common need either for IST1 itself or for the proteins it engages with (e.g. VPS4, CHMP1B, SNX15, and others). Further work will be needed to better understand the relationship between IST1 and other factors specifically involved in “slow” recycling thereby gaining insight into how IST1 contributes.

The IST1-independent “compensatory rapid recycling” that we observe aligns well with the ultrastructure of recycling tubules containing clathrin coated buds as well as TfnR visualized decades ago (Stoorvogel *et al*.). More recent reports of dynamic or “gyrating” peripheral clathrin and GGA1 (presumed to be clathrin buds on waving tubules) that are also positive for Tfn while not well understood are also consistent with the ready availability of this pathway (Zhao and Keen, 2008; Derivery *et al*., 2012; Luo *et al*., 2013; Majeed *et al*., 2014) which likely involves Rab4 marked endosomes decorated with AP-1 and GGA adaptors (D’Souza *et al*., 2014).

Finally, we found that interaction between IST1 and SNX15 on the early endosomal perimeter was direct and required the MIM1 domain of IST1. The robust expansion of an IST1/SNX15 containing subdomain when these proteins were overexpressed indicates synergy and avidity in the interactions between SNX15 and IST1. The exclusion of CHMP1B from regions coated by IST1/SNX15 – and its corresponding concentration on or near the actin-marked bases of tubules that emerge from the endosomes – raises questions for the future about how these alternating interactions are regulated.

## Materials and Methods Materials

All chemical reagents were purchased from Sigma-Aldrich unless otherwise indicated. Compounds were used at the following concentrations unless explicitly stated: 10 μM Nocodozole, 10 μM Brefeldin A. Concentrated stock solutions of all compounds were prepared in dimethyl sulfoxide (Nocodozole) or methanol (Brefeldin A) and stored at −80°C in single-use aliquots. Conjugates of Transferrin to Alexa Fluor 5565 (Cat # T35352 Invitrogen) and unlabeled Holo-Transferrin (Sigma) were used at the stated concentrations. GFP-Rab5A was a gift from Phillip Stahl (Washington University in St. Louis) and has been described in (Chen *et al*., 2014). LifeAct-mTagBFP2 was a gift from David Kast (Washington University in St. Louis) and has been described (Riedl *et al*., 2008). mEmerald-Clathrin-15 was a gift from Michael Davidson (Addgene plasmid # 54040 ; http://n2t.net/addgene:54040 ; RRID:Addgene_54040) (Fiolka *et al*., 2012). mRFP-Clc was a gift from Ari Helenius (Addgene plasmid # 14435 ; http://n2t.net/addgene:14435 ; RRID:Addgene_14435) (Tagawa *et al*., 2005). Cortactin-pmCherryC1 was a gift from Christien Merrifield (Addgene plasmid # 27676 ; http://n2t.net/addgene:27676 ; RRID:Addgene_27676) (Taylor *et al*., 2011). Apple-N1 was a gift from Michael Davidson (Addgene plasmid # 54567 ; http://n2t.net/addgene:54567 ; RRID:Addgene_54567) (Shaner *et al*., 2008). IST1 isoform A (NM_001270975.1, NP_001257904.1) and SNX15 full length or truncations as indicated were subcloned into pEGFP-N1, pcDNA4TO and mApple-N1. CHMP1B cDNA (NM_020412.4) and IST1 isoform A was recloned into pcDNA4 to create untagged CHMP1B and IST1.

### Antibodies

The following antibodies were used for immunofluorescence at the listed concentration:

| Antibody | Species | Company | Catalog # | Concentration |
| --- | --- | --- | --- | --- |
| CHMP1B | Rabbit (polyclonal) | Proteintech | 14639-1-AP | 1:200 |
| EEA1 | Mouse (monoclonal) | BD Transduction Laboratories | 610456 | 1:1000 |
| IST1 | Rabbit (polyclonal) | Proteintech | 19842-1-AP | 1:000 |
| c-myc | Mouse (monoclonal) | Developmental Studies Hybridoma Bank | 9E+10 | 1:500 |
| FK2 | Mouse (monoclonal) | Biomol/Enzo | PW8810 | 1:1000 |
| SNX15 | Mouse (monoclonal) | Santa Cruz | sc-393430 | 1:200 |
| HRS | Mouse (monoclonal) | Enzo | ALX-804-382 | 1:1000 |
| VPS35 | Rabbit (polyclonal) | Thermo Scientific |  | 1:500 |
| clathrin heavy chain (X22) | Mouse (monoclonal) | GeneTex | GTX22731 | 1:1000 |
| Cortactin (4F11) | Mouse (monoclonal) | Millipore | 05-180 | 1:250 |
| WASH | Rabbit (polyclonal) | Kind gift from Billadeau Lab MC2058 | N/A | 1:500 |
| AP-1 | Mouse (monoclonal) | Kind gift from Kornfeld Lab clone 100/3 | N/A | 1:500 |
| M6PR | Rabbit (polyclonal) | Kind gift from Kornfeld Lab | N/A | 1:1000 |
| CHMP4B | Rabbit (polyclonal) | Kind gift from Shiels Lab | N/A | 1:200 |
| TfnR | Mouse (monoclonal) | Santa Cruz | sc-32272 (3B8 2A1) | 1:500 |

Goat anti-rabbit and mouse secondary antibodies conjugated to Alexa 488, 555 or 647 were from Molecular Probes (Thermo Fisher).

### Cell culture

U2OS and T24 cells originally derived from ATCC were cultured in DMEM (U2OS) or McCoy’s (T24) (Invitrogen) with 8-10% FBS (Atlanta Biologicals). All cells were maintained at 37°C and supplemented with 5% CO2. GFPRab5B, IST1GFP, SNX15GFP stable lines were generated by placing transfected cells under selection for neomycin resistance with 500µg/mL Geneticin (Invitrogen).

### Plasmid Transfections

Cells were transfected with the indicated plasmid(s) using Lipofectamine 2000 following the manufacturer’s instructions and used for experiments within 16–24 hr. For immunostaining and live cell imaging, U2OS cells were reverse transfected (i.e. plated and transfected at the same time). Cells were plated onto glass slips (#1.5, Electron Microscopy Sciences) or 35 mm glass bottom dishes with #1.5 cover glass (Cellvis) and imaged or fixed 12-18hrs later.

### siRNA treatment

All siRNA duplexes used have been previously validated or used as indicated below. The following sequences were synthesized by Dharmacon and used at the following concentrations:

IST1 #1 (50nM) 5′-GCAAAUACGCCUUUCUCAUdTdT-3′ (Allison *et al*., 2013) IST1 #2 (40 nM) 5’GGCCUUAUCCAGUCAUAUGAdTdT-3’

CHMP1B (25nM) 5’-UGGACAAAUUCGAGCACCAdTdT-3’ (Morita *et al*., 2011)

Control (non-targeting firefly luciferase) (25nM) 5′-AUGUAUUGGCCUGUAUUAG-dTdT-3’) (Skowyra *et al*., 2018)

Control LAP1 (25nM) 5’-CGUCUUUCCUCUAGUACUA-3’ (Goodchild *et al*., 2015) Control (Silencer Select Negative Control #1 siRNA (Ambion, cat no 4390843))

CHMP4B (25nM) 5’-CGAUAAAGUUGAUGAGUUAdTdT-3’ (Morita *et al*., 2011)

Cells were plated at 90,000 (T24) or 150,000 (U2OS) per 6 well dish, and 3-5 hrs after plating cells were transfected using DharmaFECT 1 (Dharmacon) according to manufacturer’s instructions. Cells were replated after 24 hrs onto slips or live cell dishes, uncoated or coated with fibronectin (10μg/mL Fibronectin for 1hr at RT, washed 2X in PBS, then covered in media before cells added) as indicated. The cells were then imaged or fixed after 48 hrs of siRNA treatment. Alternatively, after 24 hrs IST1 siRNA treated cells were replated and transfected with a second IST1 siRNA transfection. These cells were replated onto slips after 24 hrs, and then fixed after a total of 96 hrs of siRNA treatment. When ESCRT-III siRNA treated cells were mixed with control siRNA cells before plating on slips (as indicated), immunofluorescence was used to identify ESCRT-III depleted cells.

### Immunoblotting

Proteins from whole cell lysates were separated by SDS–PAGE and transferred to nitrocellulose. Blots were incubated with primary antibodies against IST1 or TfnR followed by HRP conjugated secondary antibodies in Tris-buffered saline containing 0.1% Tween-20 and 5% milk. Proteins were detected by chemiluminescence using a ChemiDoc MP imaging system (BioRad).

### Microscopy

The following light microscope platforms were used for imaging.

| Microscope Platform | 1 Widefield fluorescence | 2 Spinning disk confocal | 3 Spinning disk confocal | 4 Spinning disk confocal | 5 Airy Scan confocal |
| --- | --- | --- | --- | --- | --- |
| Microscope | Olympus IX-81 | Olympus IX-81 (same as Platform 1) | Nikon Ti-E inverted | Nikon Eclipse Ei2 inverted | Zeiss LSM 880 |
| Objective(s) | 60X, 1.42 NA oil immersion (PLAPON) | 60X, 1.42 NA oil immersion (PLAPON Apochromat) | 60X1.4 NA oil immersion (CFI Plan Apochromat Lambda) | 60X1.4 NA and 100X 1.45 NA oil immersion (CFI Plan Apochromat Lambda) | 63X, 1.40 oil immersion Plan Apochromat |
| Light Source/Lasers | Ex-Cite 120Q (Excelitas) | Two-line (488 and 561nm) Sapphire laser module (Coherent) | Four-line (405, 488, 561, 647nm) | Four-line (405, 488, 561, 647nm) | Six-line (405, 488, 561, 647nm used) |
| Spinning disk | N/A | CSU10 (Yokogawa) | CSU-X1 (Yokogawa) | CSU-W1 (Yokogawa) | N/A |
| Camera | ORCA-Flash 2.8 scientific CMOS (Hamamatsu Photonics) | Cascade B12 (Photometrics) | Zyla 4.2-megapixel scientific CMOS camera (Andor) | ORCA-Flash 4.0 scientific CMOS (Hamamatsu Photonics) | Airy Scan Detector |
| Acquisition software | Metamorph Advanced software | mManager | Nikon Elements | Nikon Elements | Zeiss ZEN |
| Live cell imaging chamber | N/A | Tokai Hit INU series stage-top incubator | N/A | Tokai Hit STX stable top incubator | N/A |

### Tfn live cell exocytosis assay

Cells were plated on p35 dishes with 1.5 coverslip glass bottoms (MatTek) one day before imaging. 10ug/mL Alexafluor Tfn555 was added in normal cell media (8% serum in McCoys media for T24 or DMEM for U2OS). Prior to Tfn incubation, dishes were transferred to live cell imaging chamber on microscope (Platform 4). Several positions to image (221×221 µm^2^ regions) were selected across the dish. After 1hr Tfn555 media was removed. Cells were washed with 1mL of PBS followed by 2mL of chase media consisting of full media with 200ug/mL unlabeled Tfn. A z-stack (0.25um step size) was acquired for each pre-selected cell region every 2.5 minutes for 30 minutes (for T24 cells) or every 5 minutes for 1 hr (for U2OS cells).

### Tfn live cell exocytosis assay analysis

Images of individual cells were cropped into individual files. The 0min Tfn image was used to draw a mask containing a single cell. Each slice of the time series was background subtracted using a ROI from a dish region adjacent to the cell. A maximum intensity Z projection was created for each time series. A second ROI based subtraction followed by a constant background subtraction was taken to reduce signal outside of cells to zero. The Tfn intensity was then measured for each timepoint and reported as mean Tfn intensity across each cell. To normalize Tfn signals, the total Tfn intensity (integrated density) within each cell at each time point was divided by the total Tfn intensity within the cell at the 0min time point.

### Tfn Pulse Assay

We monitored the kinetics with which a labeled pulse of Tfn moves into and through the cell as optimized (Kalaidzidis *et al*., 2015). T24 cells were fed 20ug/uL Alexafluor 555 transferrin (Molecular Probes) for 30 seconds (cells on slip inverted on drop of pulse media). This incubation was done at 37°C (dry-hot bath) in serum-free pulse media containing either 1:1 HBSS and McCoy’s Media or McCoy’s Media buffered with by 25mM HEPES to maintain pH. After pulse, slips were dropped into ice-cold serum free media to wash excess Tfn and then transferred to 37°C serum free media to continue endocytosis for 1.5min (2min total). Slips were then transferred to Chase media, consisting of serum containing media with 200ug/mL unlabeled Tfn, and allowed to incubate at 37°C (in incubator) for the indicated time. Alternatively, we also used a shorter pulse by washing slips in ice cold serum free media for 1.5 minutes after the 30 second pulse, before place in chase media for indicated times. This limited the uptake to the pulse time alone. For assessing total Tfn binding, cells were incubated with labeled Tfn (inverted on a drop of pulse media) on ice for 30 min and washed in ice cold PBS before freezing. All slips were frozen in < −20°C methanol followed by fixation and immunostaining as below.

### Immunofluorescence Microscopy

Cells were fixed using using two approaches: either cross-linking fixative alone (paraformaldehyde) followed by TX-100, or precipitation fixation (-20°C methanol) followed by rehydration in cross-linking fixative (bis(sulfosuccinimidyl)suberate) as described in (Bhattacharyya *et al*., 2010). For cross-linking fixative, cells were incubated in 3% PFA, 4% sucrose for 20 minutes, and then permeabilized with either 0.1% saponin or 0.1% Triton for 10 min (as indicated). For quantification of TfnR present on cell surface, cells were fixed in 3% PFA, 4% sucrose and then immediately processed for staining without permeabilization. Methanol + BS3 fixed cells were process as described (Bhattacharyya *et al*., 2010) with the slight modification that 12mm slips were dropped in 5mL of methanol in a 15mL conical tube, precooled to −80C to ensure that the methanol was <− 20°C when slips were added. All subsequent IF steps were identical for both processes, starting with inverting the slips on blocking buffer (1hr at RT), and then in primary antibodies in block (overnight at 4°C), washing, as described in (Bhattacharyya *et al*., 2010), and then inverting in secondary antibodies in block (30min at RT), washing, and mounting in gelvatol. Fixed cells were imaged on platforms 1-5 as indicated.

### Image Analysis and Quantification of immunostained cells

ROIs were traced to define masks containing individual cells. For quantification of signal within masked area, image stacks were subjected to 50pixel rolling background subtraction (unless otherwise noted) in FIJI prior to preparing a maximum intensity projection. Max projections were further background subtracted to set cytoplasmic and/or dark noise to zero, and all images within the same experiment were treated identically. Image processing and analysis was completed with custom-written scripts in FIJI. Alternatively, max projections and masks were imported into Matlab using Bio-Formats (Linkert *et al*., 2010). Custom-written scripts, using the Matlab Image Processing Toolbox, were used for image analysis to measure intensities, channel correlations, and dispersal using K-means clustering. K-means clustering analysis was performed on the highest intensity pixels within a given cell—with the number of pixels depending on the area to be counted and the resolution of the image. The distances of each pixel from the identified K-means center for the cell was then plotted in a histogram. Equivalent binning was used for replicate cells so that the pixel count per bin could be averaged for all cells of a particular condition. Image processing scripts are available upon request. Quantification of SNX15GFP endosomes positive for endosomal markers (Figure 8E) was done using threshold-defined ROIs of SNX15GFP with the scorer assessing if each ROI contained or was immediately adjacent to other endosomal markers.

### Brefeldin A Assay

IST1 siRNA treated cells and control cells were either plated together onto glass slips the day before BFA treatment. Cells were fixed before or after incubation in 10uM BFA in normal media at the indicated times. T24 cells were stained with IST1 (to distinguish control and IST1 depleted cells) and TfnR. Tubular networks induced by BFA were assessed by a blinded scorer counting the number of continuous tubular structures longer than 12μm.

### Rab5 tubule quantification

Movies of GFP-Rab5b cells were acquired on Platform 2. 33ms exposures were collected every 2s over 30 sec. To quantify tubule length and duration, we segmented Rab5bgfp labelled endosomes in each image using a model-based segmentation algorithm (Paul *et al*., 2013) implemented in the Fiji plugin Squassh (Rizk *et al*., 2014). As described (Rizk *et al*., 2014), the segmentation mask assigns each pixel a value (0-1) based on its probability of belonging to an object and is the result of simultaneous image denoising, deconvolution, and segmentation. Using Squassh, background was subtracted from the original image with a rolling ball window of 10 pixels. Segmentation was determined using a theoretical PSF model calculated for our spinning disk microscope, a regularization parameter of 0.05, and a Poisson noise model. Using MATLAB (MathWorks), the resulting segmentation mask was converted into a binary image by thresholding to include only the top 3% of pixel values (representing the 3% of pixels most likely to belong to an object). ROIs were determined from connected thresholded regions, and only ROIs with an eccentricity of greater than 0.7 and a major axis length of greater than 20 pixels (5μm) were considered, empirically chosen to eliminate the majority of ROIs corresponding to non-tubular or short tubular endosomes. Note that this length criterion provides a conservative estimate of tubule length as tubules that are not linear (i.e. bent or curved) and are actually longer than 5μm are excluded. A blind scorer then determined if the remaining ROIs, or ‘tubule candidates’, were actual tubules (to be scored) or a cluster of endosomes (not to be scored) for every frame analyzed. See fig. S5 for further details and example of quantification.

### WGA Assay

U2OS cells were washed in ice cold PBS and inverted on drop containing 2.5ug/mL TRITC labeled WGA (2.5mg/mL) in serum-free pulse media containing 1:1 HBSS and McCoy’s Media. WGA-TRITC was allowed to bind for 30 minutes, then slips were washed in ice-cold serum free media for 1 minute. Slips were either fixed in PFA or allowed to endocytose WGA-TRITC labeled proteins at 37°C. Slips were transferred to serum free DMEM with or without 10uM nocodozole at 37°C (in incubator) for different intervals (5-15 minutes). Slips were either fixed in PFA (retaining PM associated WGA) or were washed in ice-cold PBS with 0.1M GlcNAc (Sigma) for 2 minutes and then fixed in PFA (to compete off WGA-TRITC bound to PM). Cells were then permeabilized (as described) and stained with antibodies.

### WGA Assay analysis

Cells were imaged using Platform 3. A z-stack was acquired for each image. Background subtraction (with rolling ball radius of 3 pixels or 270nm) was performed in image J for each slice. A maximum intensity projection of the stack was then used for analysis, Background intensity levels of non-cell area was confirmed to be zero, and additional background (150) was subtracted across all pixels and all channels to remove diffuse signal within the cell. WGA and EEA1 ROIs were then defined by using an additional threshold of 150, so as to exclude smaller, less bright WGA or EEA1 signal. Once these ROIs were defined they were dilated by two pixels (180nm) so as to include any signal immediately adjacent to them. The ROI-based correlation was the Pearson’s or Spearman’s coefficient between the sum intensity of two signals within the ROI, while the pixel-based correlation was the Pearson’s or Spearman’s coefficient between the intensity of the two signals within the pixels contained in the ROI. Nocodozole treatment was critical to disperse endocytosed material to allow assignment of signal within a specified region to a particular endosome. To determine if WGA or EEA1 ROIs were positive for a signal, the following procedure was used: if a WGA or EEA1 ROI contained any non-zero signal (from the background subtracted image) it was counted at EEA1, WGA, HRS, or SNX15 positive.

### Statistical Analysis

Data was graphed and statistical analysis was performed in Prism. Statistical tests were used as described.

### Transmission Electron Microscopy

T24 cells were plated onto fibronectin coated sapphire disks. Cells were high pressure frozen using a Leica EM High-Pressure Freezer (WUCCI core). Prior to immediate freezing cells were kept at 37C in 1:1 McCoy’s and HBSS to maintain pH. Samples were freeze substituted using Leica EM AF2S Freeze Substitution System (WUCCI core) and subsequently sectioned and imaged using the TEM described above.

## Acknowledgements

Imaging experiments were performed in part through the use of Washington University Center for Cellular Imaging (WUCCI) supported by Washington University School of Medicine, The Children’s Discovery Institute of Washington University and St. Louis Children’s Hospital (CDI-CORE-2015-505 and CDI-CORE-2019-813) and the Foundation for Barnes-Jewish Hospital (3770 and 4642). Confocal/super-resolution data was generated on a Zeiss LSM 880 Airyscan Confocal Microscope which was purchased with support from the Office of Research Infrastructure Programs (ORIP), a part of the NIH Office of the Director under grant OD021629. This research was supported by NIH grants R01GM076686 (P.I.H.) R01GM136925 (D.J.K), R01GM138448 (S.J.) and NSF fellowship DGE-1143954 (A.K.C.).

