## Supplemental Figures for "IST1 regulates select recycling pathways"

### Supplemental Figure 1

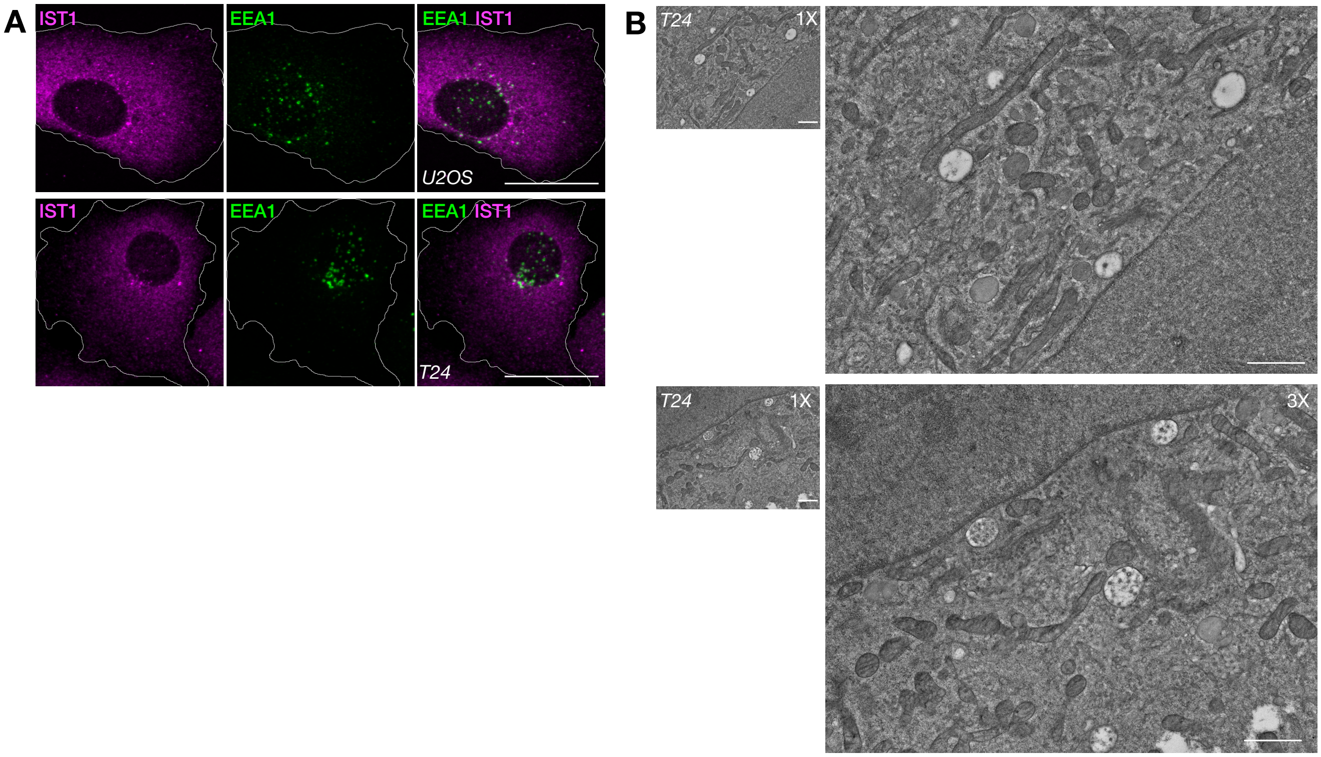


Fig. S1: **T24 cells have large endosomes. (A)** *Individual channels of whole cell images shown in Figure 1B.* **(B)** *TEM of high-pressure frozen and freeze-substituted cells T24 cells. Smaller image (1X) is at same magnification as insets in (Figure 1B). Scale bars in (A) are 25μm, and scale bars in (B) are 1μm.*

Supplemental Figure 2

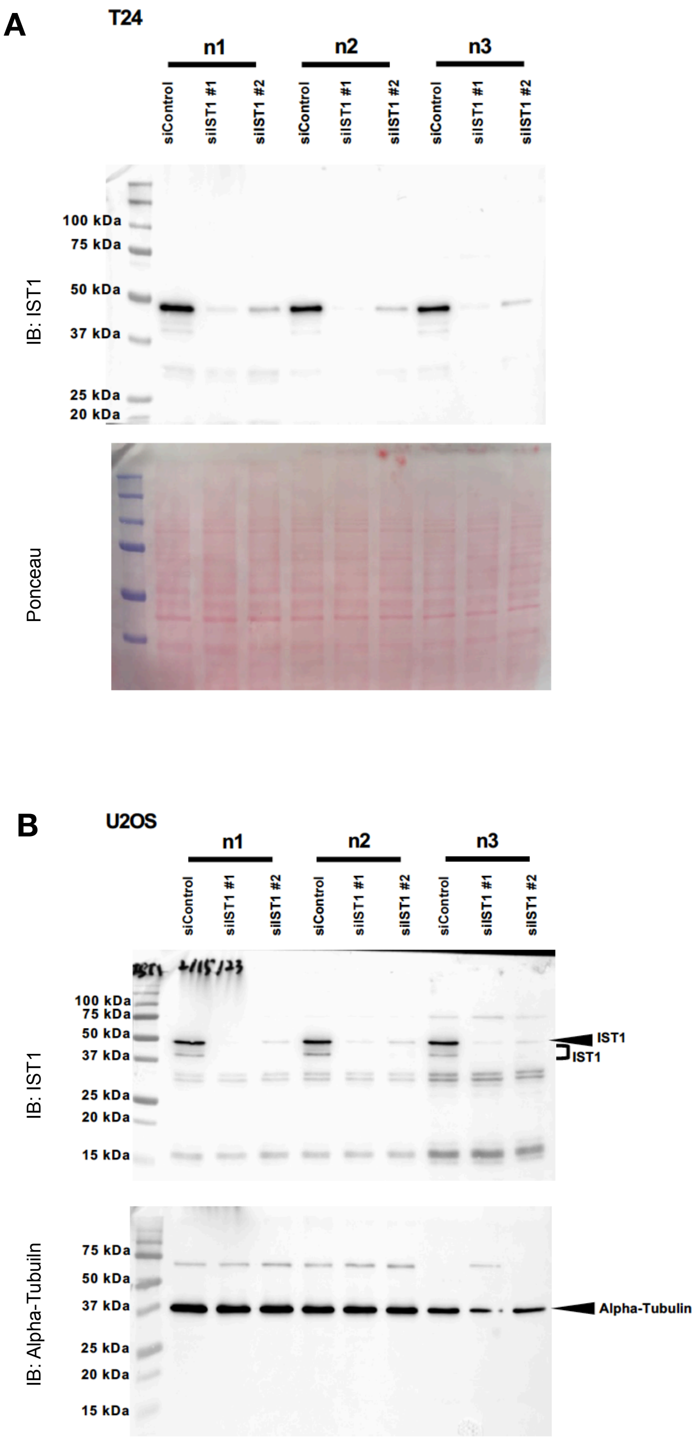


<u>Fig. S2</u>: ***Knockdown efficiency of IST1 in T24 and U2OS cells.*** *Three independent experiments of T24* ***(A)*** *and U2OS* ***(B)*** *cells treated with previously validated IST1 siRNA (siIST1 #1), a second IST1 siRNA (siIST1 #2) or control siRNA (siControl) and immunoblotted for IST1 (top panels). Loading controls are shown in bottom panels using Ponceau stain (A) or immunoblotting for alpha-tubulin (B).*

Supplemental Figure 3

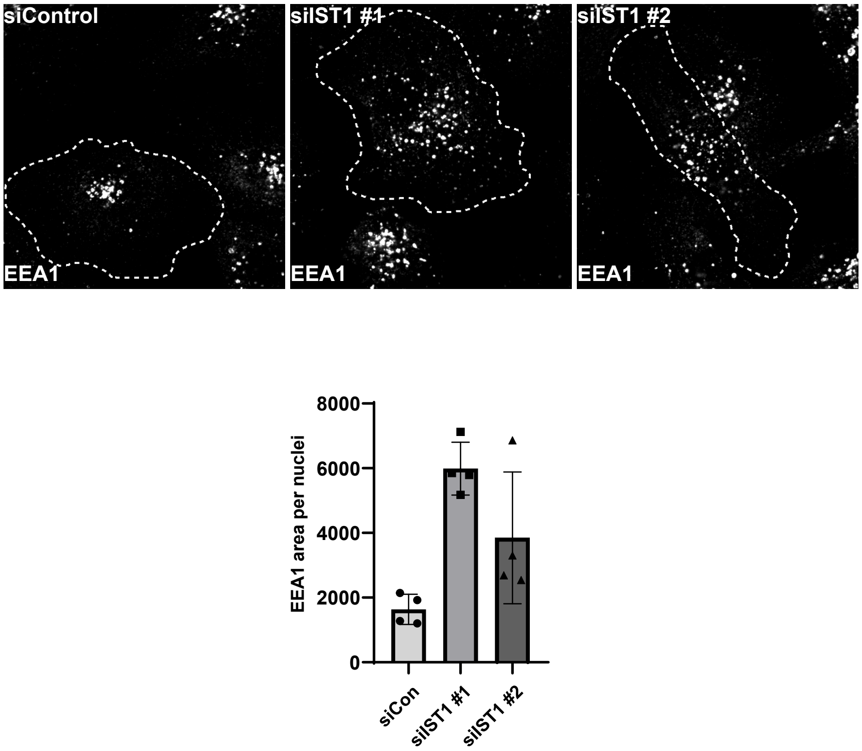


<u>Fig. S3:</u> ***IST1 depletion increases EEA1 area.*** *Cells treated with control, previously validated IST1 siRNA (siIST1 #1), or a second IST1 siRNA (siIST1 #2) were stained for EEA1. EEA1 on endosomes was quantitated as the total area above a thresholded EEA1 intensity. Each point represents average EEA1 area for a field of cells (containing > 5 cells) normalized by the number of nuclei (i.e. average EEA1 area per cell).*

Supplemental Figure 4

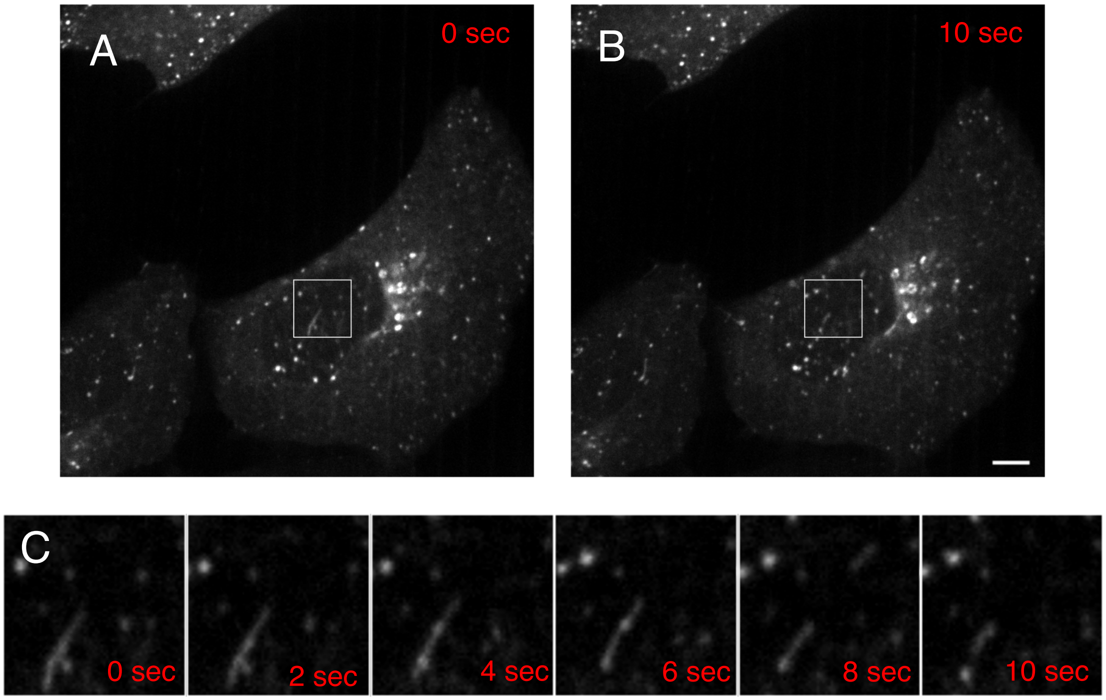


*<u>Fig. S4:</u>* ***GFP-Rab5B compartment dynamics**. Still images captured in an example time series monitoring GFP-Rab5b endosome morphology.* ***(A and B)*** *The entire frame at 0 and 10 seconds time points, respectively.* ***(C)*** *Boxed region enlarged and shown at 2 second intervals. Note apparent fission events between 6 and 8 seconds, and also between 8 and 10 seconds. Spinning disc confocal imaging, scale bar = 10 µm.*

### Supplemental Figure 5

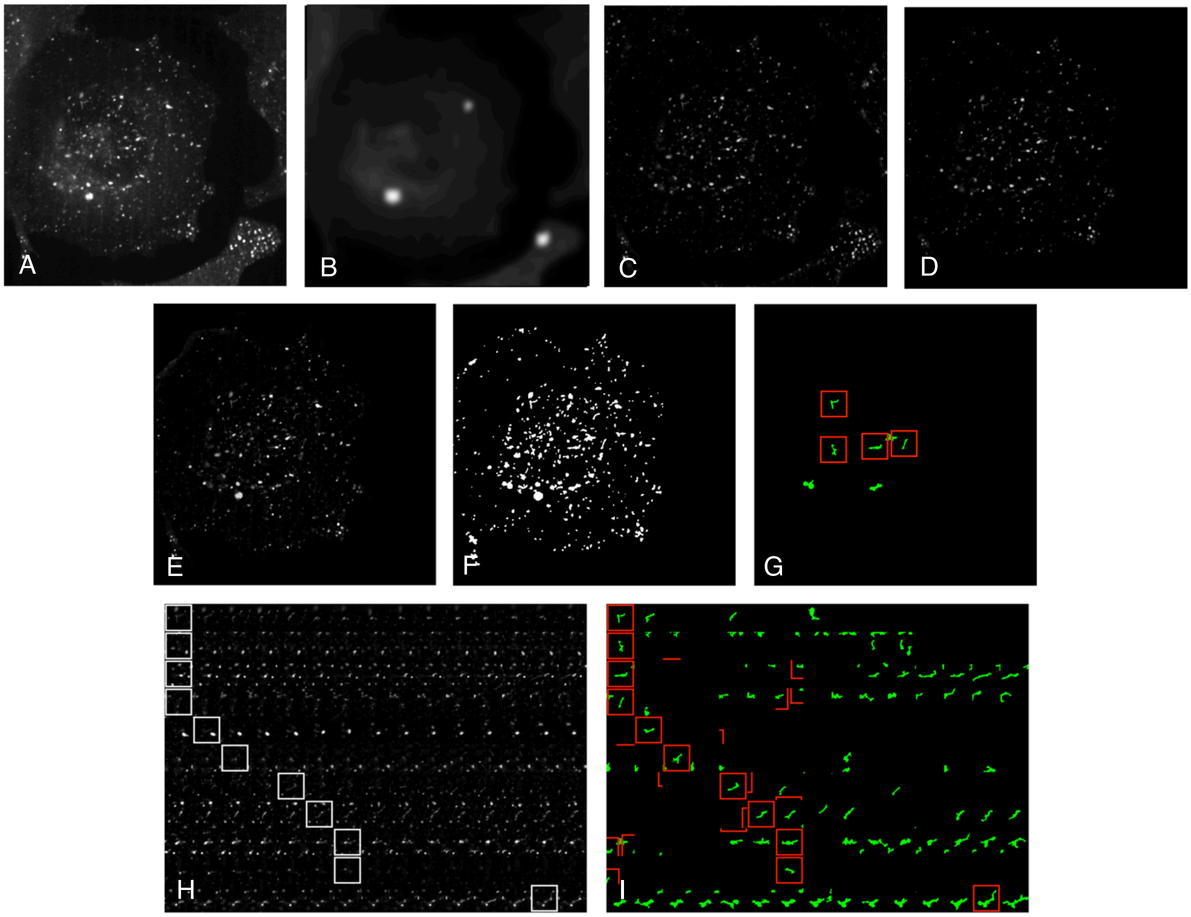


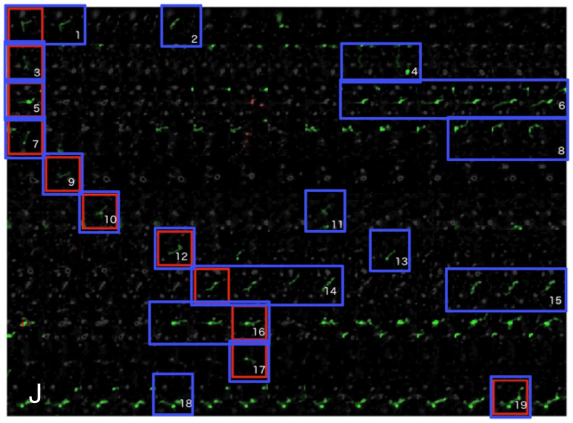


K

| Tubule # | Duration |
|---|---|
| 1 | 2 frames |
| 2 | 1 frame |
| 3 | 1 frame |
| 4 | 2 frames |
| 5 | 1 frame |
| 6 | 6 frames |
| 7 | 1 frame |
| 8 | 3 frames |
| 9 | 1 frame |
| 10 | 1 frame |
| 11 | 1 frame |
| 12 | 1 frame |
| 13 | 1 frame |
| 14 | 4 frames |
| 15 | 3 frames |
| 16 | 3 frames |
| 17 | 1 frame |
| 18 | 1 frame |
| 19 | 1 frame |

L

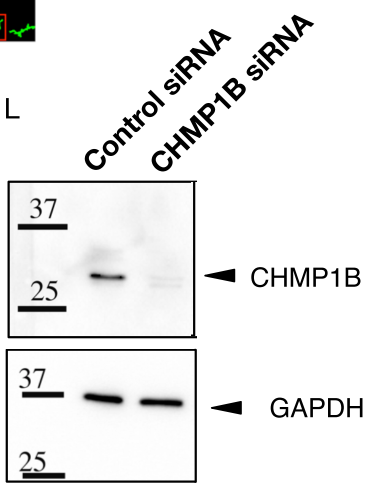

*Fig. S5:* ***Analysis of GFP-Rab5b compartment dynamics.*** *Movies collected for analysis were each a total of 15 frames, with a frame taken every 2 seconds.* ***(A-G)*** *Image processing and analysis done for each frame of a movie:* ***(A)*** *the original image,* ***(B)*** *the background determined with a rolling ball window size of 10 pixels,* ***(C)*** *the background subtracted image (B subtracted from A),* ***(D)*** *image cropped to contain single cell,* ***(E)*** *the image model produced from the SQUASSH workflow, range of pixel values is 0 to 1,* ***(F)*** *a binary image created by setting the threshold to include the top 3% of pixel values,* ***(G)*** *'Tubule candidates' or ROIs with an eccentricity greater than 0.7 and a major axis length greater than 20 pixels (5µm) shown in green. A blind scorer determined if a 'tubule candidate' was an actual tubule (boxed in red, to be scored) or a cluster of endosomes (not to be scored).* ***(H-J)*** *show results of A-G for all frames of a movie. The first 4 rows of the montage show the areas of the boxed tubules identified in G, and each column is each area for a single frame of the movie.* ***(H)*** *The same montage from the SQUASSH image model.* ***(I)*** *The montage from the green 'tubule candidate' ROIs. For final scoring, H and I were overlayed to produced (J) and identify 'tubule candidates' (any object that appeared green) which could then either be ignored or counted as a tubule by a blinded scorer.* ***(J)*** *Blue boxes outline the number of frames that a tubule 5µm or longer remained 5µm or longer.* ***(K)*** *Table shows the number of frames each tubule (labeled in J) remained 5 µm or longer. Note that any tubules lasting for a single frame (i.e. less than 2 seconds) were not counted in the final quantification in Fig. 1.* ***(L)*** *Cell shown in (A) is treated with CHMP1B siRNA, and western blot demonstrates degree of depletion of CHMP1B in U2OS cells.*

Supplemental Figure 6

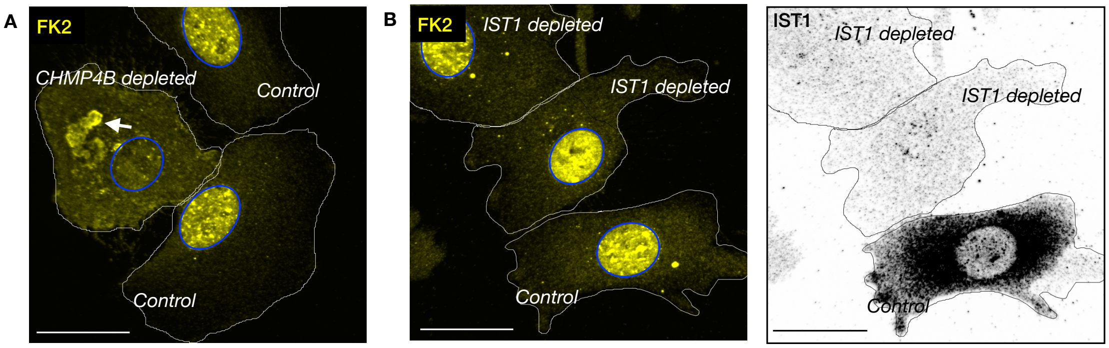


<u>Fig. S6:</u> **IST1 depletion does not lead to FK2 accumulation.** *(A-B) T24 cells individually treated with control or indicated ESCRT-III siRNA and then mixed and plated together and stained for FK2 (shown in A) as well as CHMP4B (not shown) or IST1 (shown in B). The cell nuclei are outlined in blue, and FK2 shows redistribution from nuclear signal to cytoplasmic inclusions (arrow) in CHMP4B depleted cells, not IST1 depleted cells. Scale bars are 25μm.*

Supplemental Figure 7

A

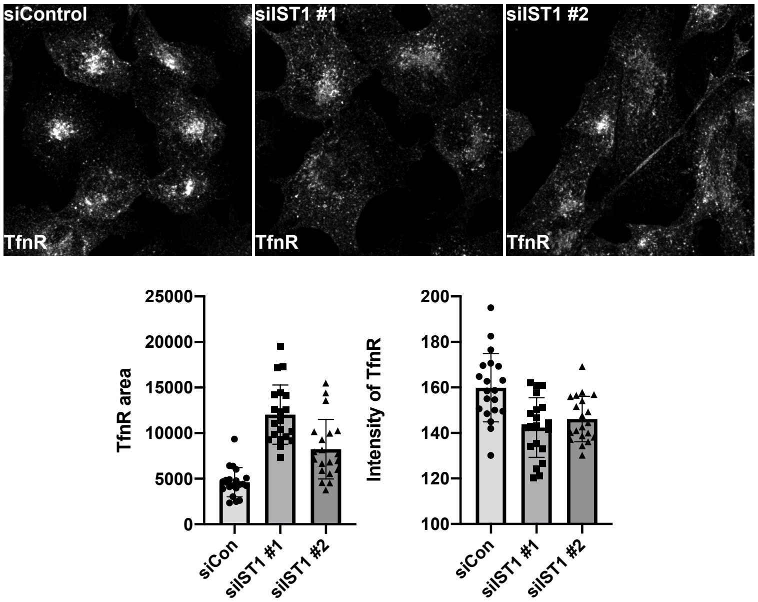


B Tfn surface binding 30 min at 4°C

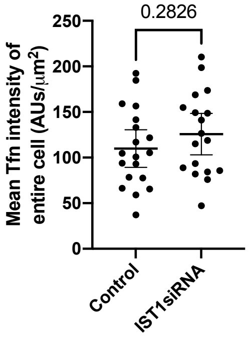


***<u>Fig. S7:</u> Depletion of IST1 leads to dispersal of steady-state TfnR from juxtanuclear localization, and no difference in amount of surface TfnR that can bind Tfn. (A)*** *Cells treated with control, previously validated IST1 siRNA (siIST1 #1), or a second IST1 siRNA (siIST1 #2) were stained for TfnR. TfnR area was quantitated as the total area of thresholded TfnR intensity, while TfnR intensity is the total (unthresholded) TfnR signal (n = 20 cells for each group).* ***(B)*** *Control or IST1 siRNA treated cells were allowed to bind labeled Tfn555 on ice for 30min and were immediately fixed. Graph shows mean±95%CI. Control and IST1 siRNA treated cells n= 19, n=18 respectively. Statistical analysis using student's t test showed no significant difference in PM Tfn levels in IST1 depleted results. This lack of a significant difference was replicated in two more independent experiments.*

Supplemental Figure 8

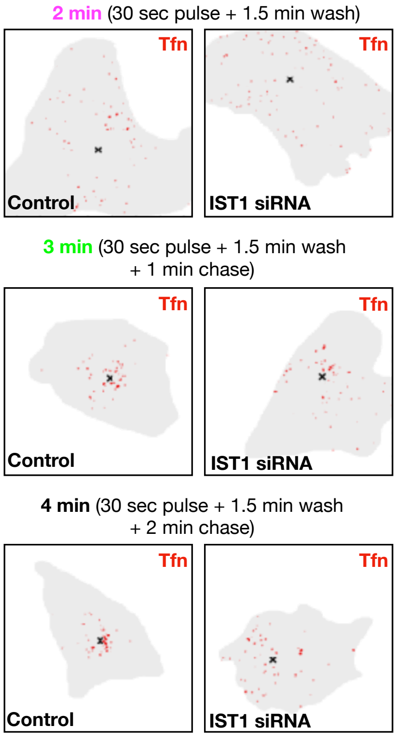


*Fig. S8*: ***Loss of IST1 impairs the trafficking of endocytosed cargo, as assessed by k-means clustering.*** Cell masks are shown in grey, and red pixels show the locations of the highest Tfn pixel intensities (within a total area of 4$\mu m^2$) in the cell. The X denotes the center identified by k-means clustering of the Tfn areas. This intensity-independent assay enables a quantitative way to detect dispersed endocytosed Tfn after 2min of endocytosis and the efficiency of cargo clustering after 4 min.

Supplemental Figure 9

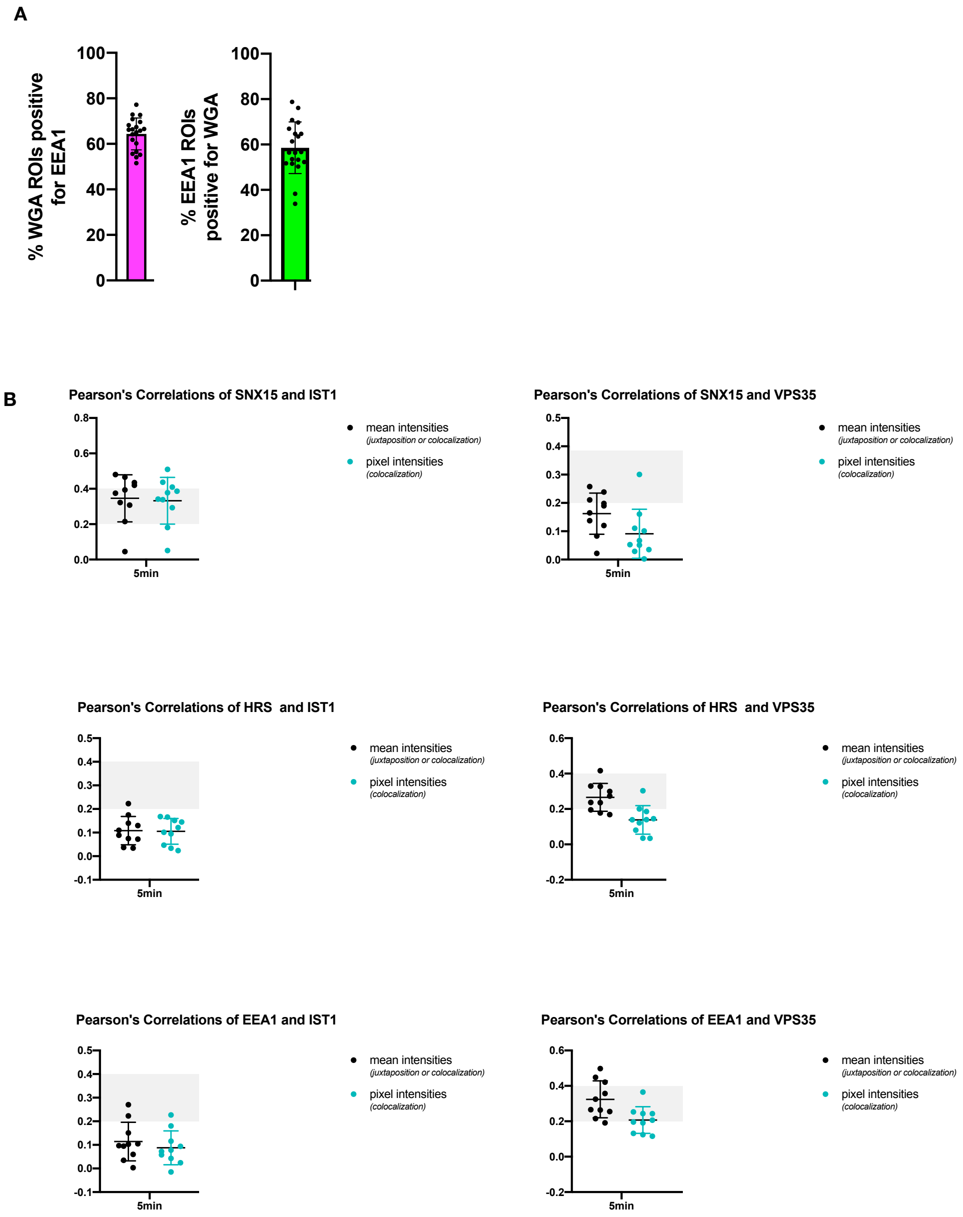


Fig. S9: ***A majority of internalized WGA localizes to EEA1 sorting endosomes, while SNX15 co-localizes and VPS35 co-distributes with IST1. (A****) Percent of WGA ROIs positive for EEA1 within cells, and percent of EEA1 ROIs positive for WGA (n=20).* ***(B)*** *Pixel-based and ROI-based Pearson's correlations for IST1 and VPS35 co-stained with either SNX15, HRS, or EEA1 in WGA assay.*

Supplemental Figure 10

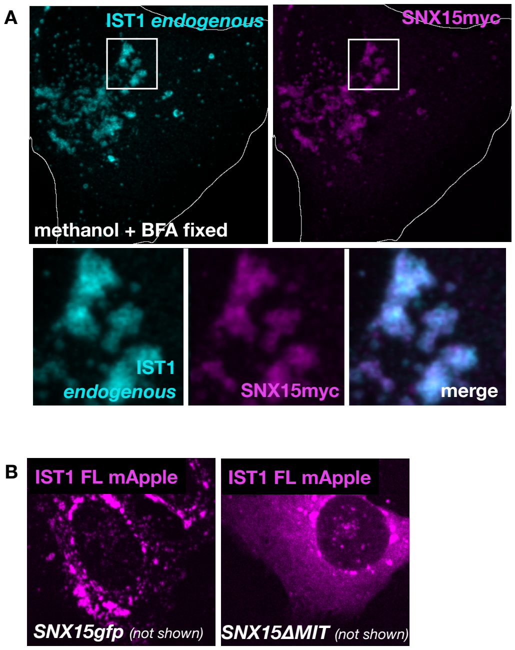


<u>*Fig S10.*</u> ***IST1 recruitment to endosomes is promoted via the MIT domain of SNX15***
**(A)** U2OS cells expressing exogenous SNX15myc were harvested, stained for endogenous IST1. Cell was frozen in methanol and fixed in BS3 (see methods). (B) *Fixed U2OS cells expressing IST1mApple and SNX15GFP or SNX15ΔMIT.*

Supplemental Figure 11

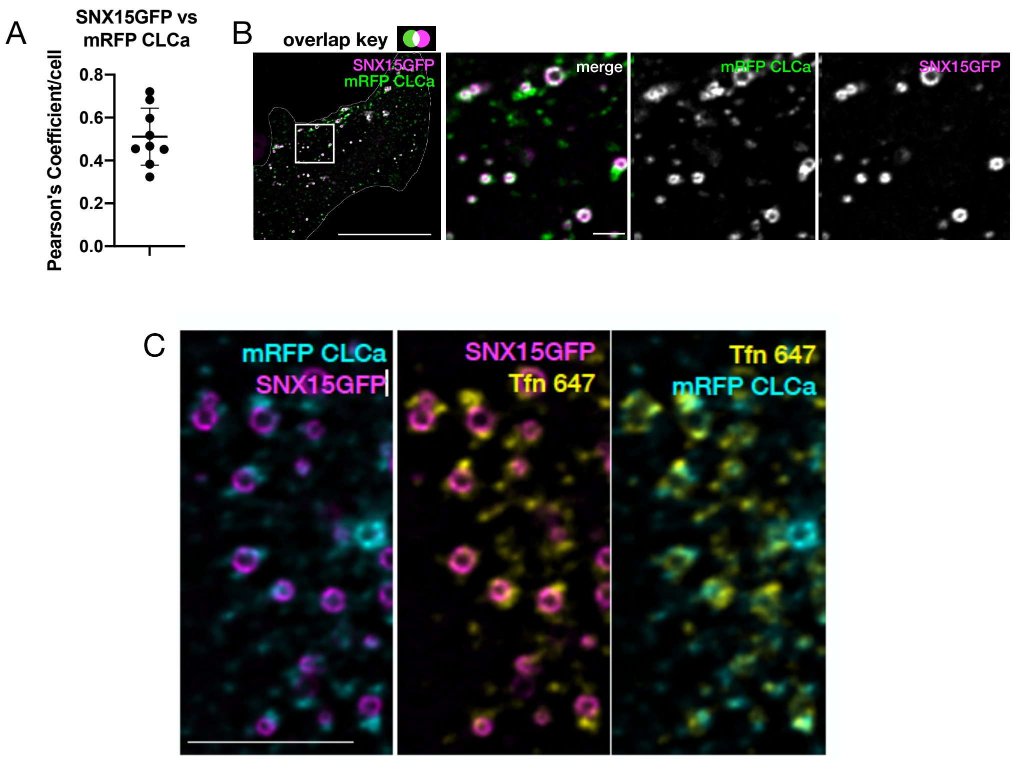


*Fig. S11:* ***Live cell imaging of SNX15GFP and CLCa****.* ***(A)*** *Quantification of the Pearson's correlation coefficient in cells expressing SNX15GFP and mRFP CLCa, which includes cell in Figure 9A and cell in* ***(B)****. Cell shown in Figure 9A had a coefficient on low end (0.45), while cell shown in (B) has coefficient on high end (0.72).* ***(C)*** *Cells over-expressing SNX15GFP and mRFP CLCa allowed to endocytosed Tfn647 for 1 hr. Some Tfn positive tubules on SNX15gfp endosomes are positive for clathrin. Scale bars in (B) are 25µm (cell) and 1µm (inset), and scale bar in (C) is 5µm.*
